# Tumor-associated stroma shapes the spatial tumor immune microenvironment of primary Ewing sarcomas

**DOI:** 10.1101/2025.01.31.635996

**Authors:** Christopher Kuo, Krinio Giannikou, Nuoya Wang, Mikako Warren, Andrew Goodspeed, Nick Shillingford, Masanori Hayashi, Micha Sam Brickman Raredon, James F. Amatruda

## Abstract

To date, few studies have detailed the tumor microenvironment (TME) of Ewing sarcoma (EwS). The TME has a vital role in cancer survival and progression with implications in drug resistance and immune escape. By performing spatially resolved transcriptomic analysis of primary treatment-naïve EwS samples, we discovered greater stromal enrichment in localized EwS tumors compared to metastatic EwS tumors. Through spatial ligand-receptor analysis, we show that the stromal enriched regions harbor unique extracellular matrix related cytokines, immune recruitment and proinflammatory microenvironmental signals, implying EwS stroma may play an anti-tumorigenic role by acting as an immune recruitment center. All EwS tumors expressed pro-tumorigenic MIF-CD74 immune signaling, suggesting a potential immune-evasive mechanism and immunotherapy target. Our findings provide insight into tumor cell/stromal cell interactions in EwS and serve as a valuable resource for further investigations in the tumor immune microenvironment of EwS.

## Introduction

Ewing sarcoma (EwS) is a malignant bone and soft-tissue tumor that occurs most commonly in children, adolescents and young adults.^1,2^ Less than 30% of patients with initial metastases will survive for 5 years.^3,4^ New therapeutic approaches are needed given the vast majority of children with metastatic or relapsed EwS die despite intensive, multimodal therapy.^5^ Additionally, survivors face a lifelong risk of adverse health effects due to toxicity of therapy, including secondary cancers. EwS is caused by chromosomal translocations between a FET family member and an *ETS*-type gene, the most common of which is a t(11;22) that creates the *EWSR1::FLI1* fusion oncogene.^6^ Despite the discovery of the *EWSR1::FLI* rearrangement more than 30 years ago, effective targeted therapies remain unavailable.^7^ Over the past decade, immunotherapy has revolutionized the field of cancer therapy. Instead of directly targeting cancer cells, immunotherapy harnesses the host’s own immune system to eliminate cancer cells.^8^ Immunotherapy such as checkpoint inhibition through programmed death 1 (PD-1) blockade has recently achieved complete response in all patients with mismatch repair-deficient, locally advanced rectal cancer.^9,10^ Similar effectiveness with immunotherapy is seen with CAR-T therapies in leukemias.^11^ However, despite advances in our understanding of immunobiology and cancer genomics, immunotherapy has not been successful in EwS.^12^ While low tumor mutational burden and resulting scarcity of tumor infiltrating lymphocytes have been proposed as an explanation for the lack of immunogenicity of EwS, the potential immunosuppressive roles of the innate immune system and the tumor microenvironment (TME) in EwS have not been fully elucidated.^6^

Previous studies investigating the EwS TME have identified macrophages through conventional immunohistochemistry (IHC) using CD163 or CD68 as single-plex marker of macrophages.^13,14^ However, while informative, conventional IHC does not fully reflect the complexity of the TME, and although bulk tumor genomics has identified immunosuppressive M2 macrophages being the predominant immune cell type in EwS family of tumors,^15^ there is a paucity of literature on the spatial distribution of these immune cells within the TME. In recent years, single-cell RNA sequencing (scRNA-seq) has allowed more granular dissection of the TME of many cancer types by profiling tumor and immune cells. Visser and colleagues performed scRNAseq of 18 primary EwS samples and reported dysfunctional antigen-presenting cells as well as immunosuppressive microenvironmental signals.^16^ Cillo et al. performed scRNAseq on immune cells from blood and tumors from patients with EwS, and identified distinct myeloid cells as well as CD8+ effector T cells and exhausted CD8+ cells.^17^ Although scRNA-seq analysis resolved the composition of cell types within the TME of EwS, detailed spatial transcriptomic analysis of the TME of EwS is lacking. The intricate interactions between the host’s innate/adaptive immune system and the cancer cells lead to distinct spatial phenotypic and architectural changes within the tumor.^18^ Being able to visualize the spatial heterogeneity of cell types and cell states in intact tumor samples can provide insight into how the host’s innate and adaptive immune cells contribute to the TME. Additionally, this information could help identify novel treatment approaches to prevent tumor progression and/or invasion.^18^

Here we characterize the TME of EwS through spatial transcriptomics (ST) of primary biopsied human EwS tumors. We implemented a comprehensive pipeline for ST analysis of EwS, and noted significant enrichment of extracellular matrix-related (ECM-r) genes in primary tumors from patients with localized disease. These ECM-r genes colocalized with the tumor stroma. Gene composite analysis demonstrated that macrophage/monocyte and cancer-associated fibroblast (CAF) genes colocalized with the ECM-r genes. Fusion-driven sarcomas such as EwS have been noted to have low immune infiltrations compared to sarcomas with complex genomes.^19^ Despite this, our studies showed infiltrative T and B cells in EwS tumors. We found that within the tumor stroma there were significant ECM and immune related microenvironmental signals, indicating interactions between different types of immune cells and between immune cells and stromal cells. Tumors from patients with localized EwS (loc-EwS) showed enrichment of *TGFB* signaling in the stroma compared to tumors from patients with metastatic EwS (met-EwS). Tumor stroma harbored unique spatial microenvironmental connectivities related to tumor inhibition, immune activation and recruitment, which were predominantly observed loc-EwS tumors. Notably, we identified *MIF-CD74* immune signaling in all tumors that may contribute to a pro-tumoral and immune-evasive TME. We validated key TME features identified in our ST analysis via high-plex proteomics immunostaining at single-cell resolution. Collectively, our results provide novel insights into the spatial architecture and immune landscape of EwS tumors, suggesting possible anti-tumorigenic roles of stroma and potential mechanisms by which EwS manipulates the surrounding immune microenvironment to ultimately evade immune surveillance.

## Results

### A pipeline for spatial transcriptomic analysis of Ewing sarcoma

To define the spatial transcriptional landscape of Ewing sarcoma (EwS) tumors, we implemented a comprehensive pipeline for ST analysis of clinical tumor specimens (Figure 1). We selected formalin-fixed, paraffin-embedded biopsy tissue samples from 16 primary EwS tumors from patients who had either localized EwS or metastatic EwS at the time of diagnosis. All of the tumors were histopathologically confirmed by an expert pathologist to be Ewing sarcoma, and molecular diagnostics confirmed the presence of a *FET-ETS* fusion gene (Table 1). Quality control of the tumor specimens was performed and regions that were annotated as non-necrotic and tumor dense were then sectioned and placed on Visium spatial gene expression slides.^20^ Following sequencing, data were aligned to the human reference genome (hg19) and spatially mapped. Expression data were then analyzed via dimensionality reduction (Uniform Manifold Approximation and Projection (UMAP)), clustering and visualization through STUtility and Seurat R packages.^20,21^ Expression data for each sample were normalized using LogNormalize to optimize preservation of biological variation. After initial data processing, we assessed expression of gene markers specific to Ewing sarcoma (NKX2-2, CD99) to identify the spatial location of tumor cells.

**Figure 1:**
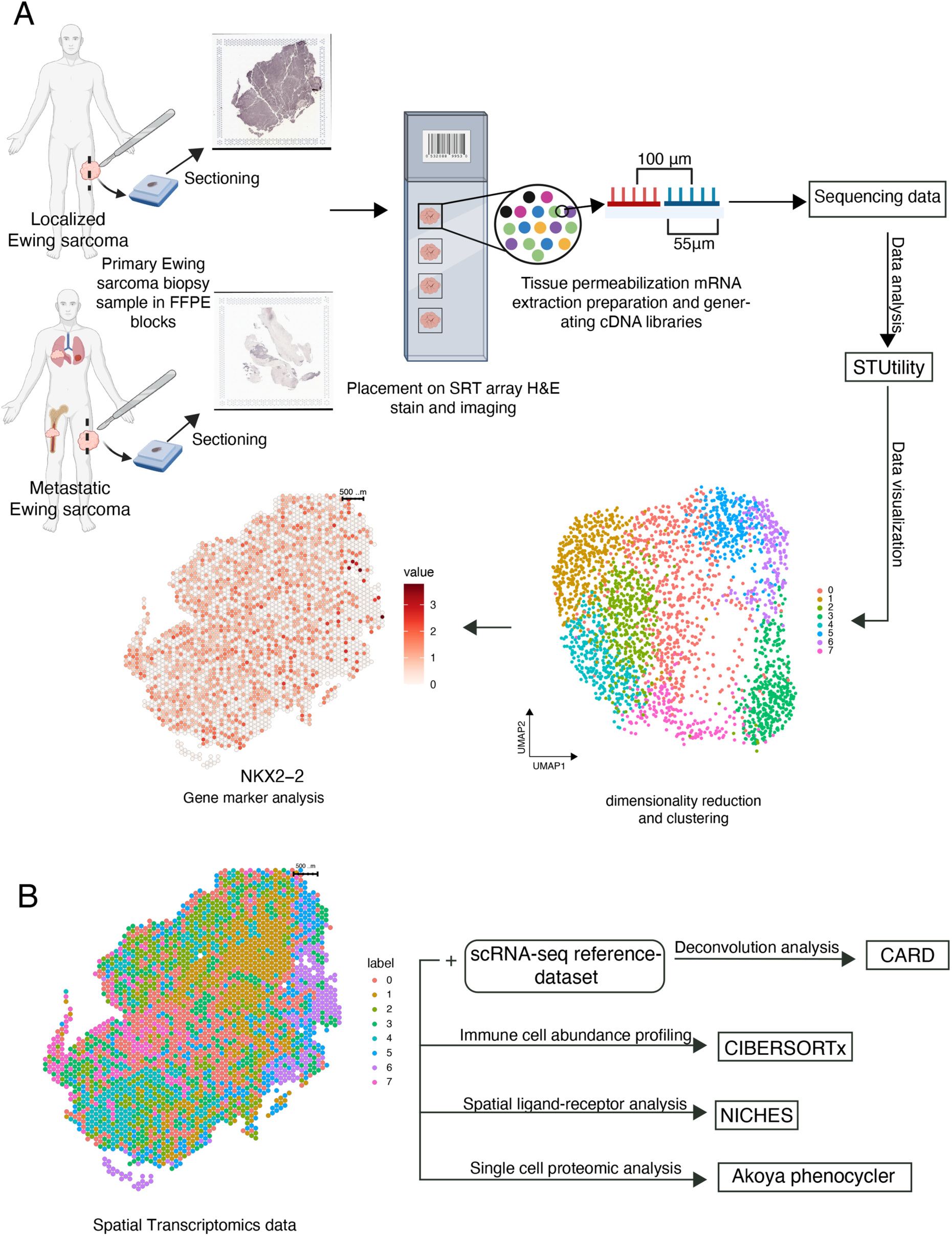
Overview of comprehensive Spatial Transcriptomic analysis. (A) Overview of experimental method, beginning with selection of biopsied formalin-fixed, paraffin-embedded primary tumor specimens from patients with localized Ewing sarcoma (EwS) and metastatic EwS. Tumor dense region was selected by an expert pathologist followed by fixation of tumor to Visium spatial gene expression slide followed by sequencing and data analysis. (B). Downstream analyses of ST datasets to further elucidate immune cell populations and spatial interactions.

We used a publicly available scRNA-seq EwS dataset as a reference to perform conditional autoregressive-based deconvolution analysis to enhance the ST resolution and infer the composition and abundance of all different cell types within each spot.^22,23^ We also profiled the immune cell abundance in all 16 tumors through CIBERSORTx.^24^ To uncover the cellular microenvironment signals from ST data, we utilized Niches Interactions and Cellular Heterogeneity in Extracellular Signaling (NICHES).^25^ Lastly, we validated the transcriptomic findings at the proteomics level by performing single cell spatial proteomics analysis using PhenoCycler.^26^ In conclusion, our ST pipeline allowed us to understand the tumor microenvironment (TME) of EwS at the spatial level.

### Integrated clustering reveals intertumoral heterogeneity of Ewing sarcoma

To further characterize the transcriptional heterogeneity between tumors, we merged and integrated ST data from all 16 tumors. When analyzed individually, each tumor had distinct clusters on UMAP plots, reflecting unique spatial transcriptional profiles (Figure 2A). Upon merging the datasets, tumors retained their distinct clustering patterns, occupying separate regions on the integrated UMAP (Figure 2B). For integration, we normalized and identified variable features for each tumor independently, and selected features that are repeatedly variable across all tumors. We applied a resolution of 0.4 to avoid overclustering of the data and to preserve biological heterogeneity. Clustering of all 16 integrated tumors produced 10 clusters, and integrated ST data from all 16 tumors overlapped on the UMAP (Figure 2C). When visualizing the integrated clusters spatially in individual tumors, each tumor had a distinct clustering pattern and had different clusters represented (Figure 2D). These findings highlight the inherent intertumoral heterogeneity of Ewing sarcoma at transcriptional spatial level, underscoring the diverse molecular landscape across tumors.

**Figure 2:**
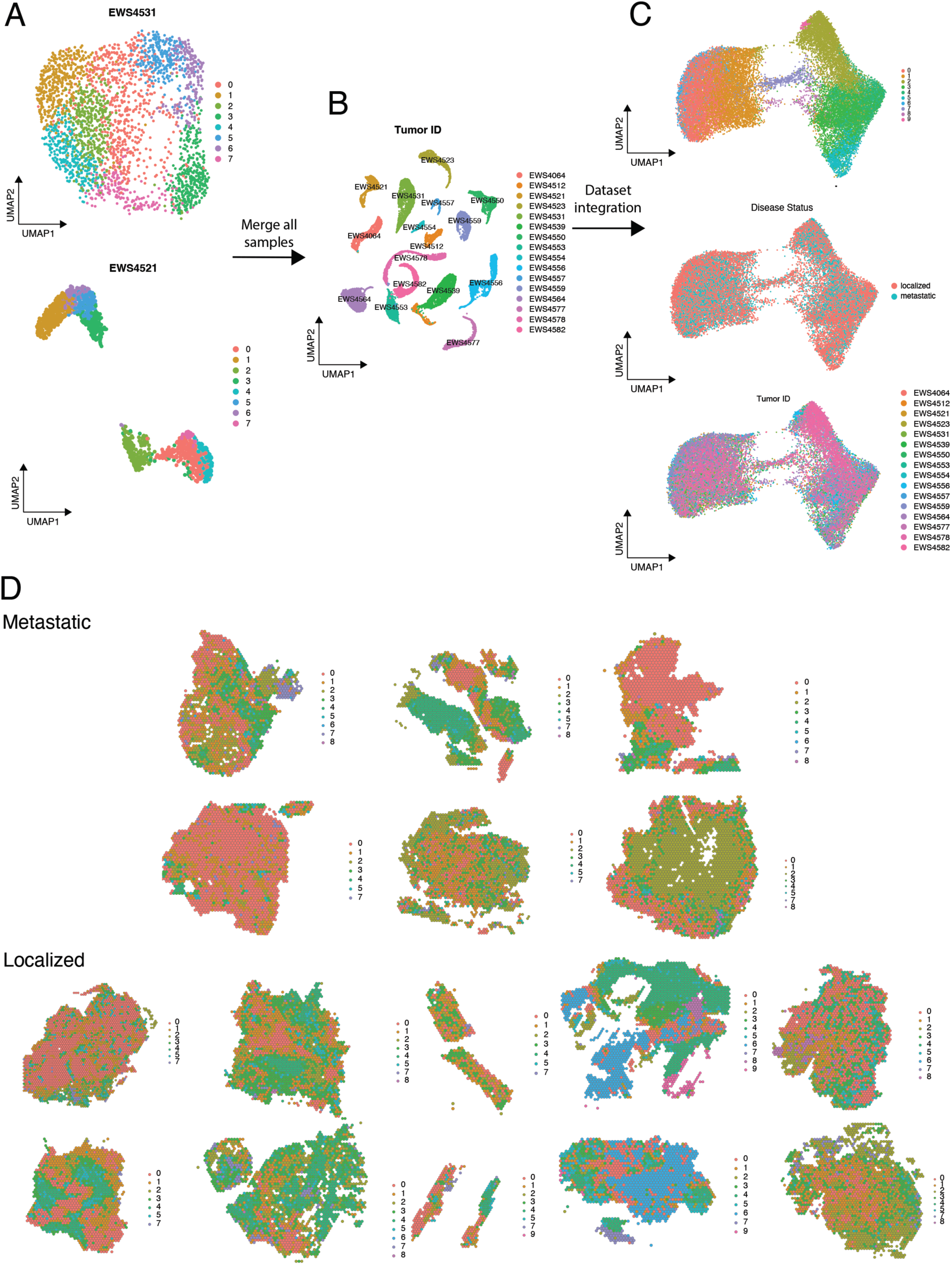
Integrated analysis of all EwS tumors reveals intertumoral transcriptional heterogeneity. (A) Representative Uniform Manifold Approximation and Projections (UMAPs) from two tumors. (B) UMAP of all 16 tumors pre-data integration. (C) UMAP of integrated 16 tumors. UMAP split by unique clustering of integrated dataset, disease status (localized vs. metastatic at diagnosis), and individual tumor samples. (D) Spatial mapping of clusters in primary tumors from select patients with metastatic or localized disease at diagnosis.

### Inference of tumor histology from gene expression

Our finding that transcriptional clustering revealed intertumoral heterogeneity of EwS led us to examine whether tumor histology could be inferred from ST gene expression data. To achieve this, we implement ‘Estimation of Stromal and Immune cells in Malignant tumours using Expression data’ (ESTIMATE) algorithm and using gene expression signatures we inferred the proportions of stromal and immune cells in tumor samples.^27^ ESTIMATE performs single-sample gene set-enrichment analysis ^28^ by leveraging curated ‘stromal gene signatures’ to capture stromal cells in tumor tissue and ‘immune gene signature’ to identify immune cells in tumor tissues.^27^ A stromal and immune scores were then generated and combined to infer a tumor purity score. Prior to ST analysis, all EwS tumors were subjected to hematoxylin and eosin (H&E) staining and digital images were reviewed by an expert anatomic pathologist. Tumor and stromal regions were annotated in blinded fashion. We then compared the annotated H&E-stained tumor tissues to gene-signature enriched annotated regions. All tumors demonstrated good agreement between histopathologic annotation and ST delineation of stromal and tumor regions (Figure 3 and Supplementary Figure S1A-D). Moreover, regions that were transcriptionally annotated as stroma overlapped with immune-rich with significant correlation (Figure S1E). Thus, these ST gene expression data recapitulated the histopathology of primary EwS tumors, reinforcing the utility of ST in capturing the complex spatial architecture of EwS.

**Figure 3:**
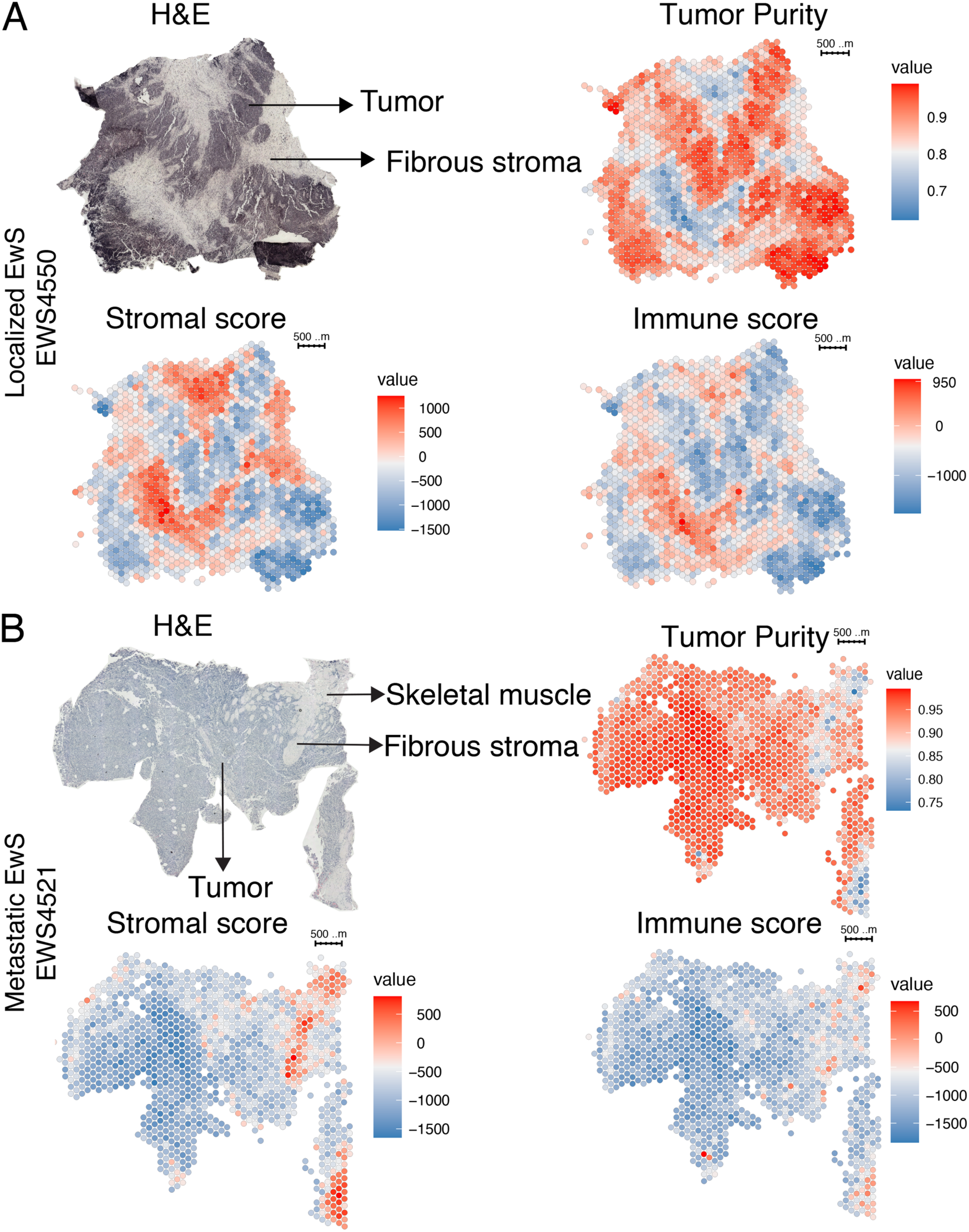
Transcriptomic annotation correlates with tumor histology. (A) Representative H&E-stained image from tumor EWS4550 (from a patient with localized disease) with tumor region and fibrous stroma region as noted by an expert pathologist. Tumor purity and stromal scores generated by ESTIMATE transcriptional analysis correlate with H&E stained images. The ESTIMATE immune score is also shown. (B) Representative H&E-stained image from tumor EWS4521 (from patient with metastatic disease) with tumor, skeletal muscle and fibrous stroma region noted by an expert pathologist. ESTIMATE analysis of EWS4521 recapitulating H&E-stained image.

### Extracellular matrix-related genes are enriched in EwS tumors from patients with localized disease

To gain further insight in the transcriptional differences between primary tumors from patients with localized EwS (loc-EwS) and primary tumors from patients with metastatic EwS (met-EwS), we merged all 16 tumors together (Figure 2B) and applied Gene Set Enrichment Analysis (GSEA) to assess differential enrichment of molecular pathways using Hallmark gene sets between loc-EwS and met-EwS. Loc-EwS tumors were significantly enriched in the epithelial to mesenchymal transition (EMT) pathway (Figure 4A).^29^ Several inflammatory molecular pathway signatures were also highly enriched in loc-EwS, such as tumor necrosis factor alpha (TNFa) signaling, interferon (IFN) alpha and IFN gamma response (Figure 4A). There was also enrichment of PI3K/AKT/mTOR signaling and transforming growth factor beta (TGF-b) signaling in loc-EwS (Figure 4A). In contrast, met-EwS exhibited enrichment in proliferative pathways such as E2F targets, MYC targets, G2M checkpoint (Figure 4A). Notably, met-EwS also exhibited downregulation of inflammatory response pathways compared to loc-EwS. Annexin A1 (*ANXA1*) was significantly overexpressed in loc-EwS compared to met-EwS. *ANXA1* was recently reported to have prognostic significance with low expression level associated with lower overall survival in both primary and metastatic EwS.^30^ We also noted upregulation of the six-transmembrane epithelial antigen of the prostate 1 (*STEAP1*) in loc-EwS tumors. High STEAP1 expression has been correlated with improved overall survival (OS) in patients with EwS.^31,32^

**Figure 4:**
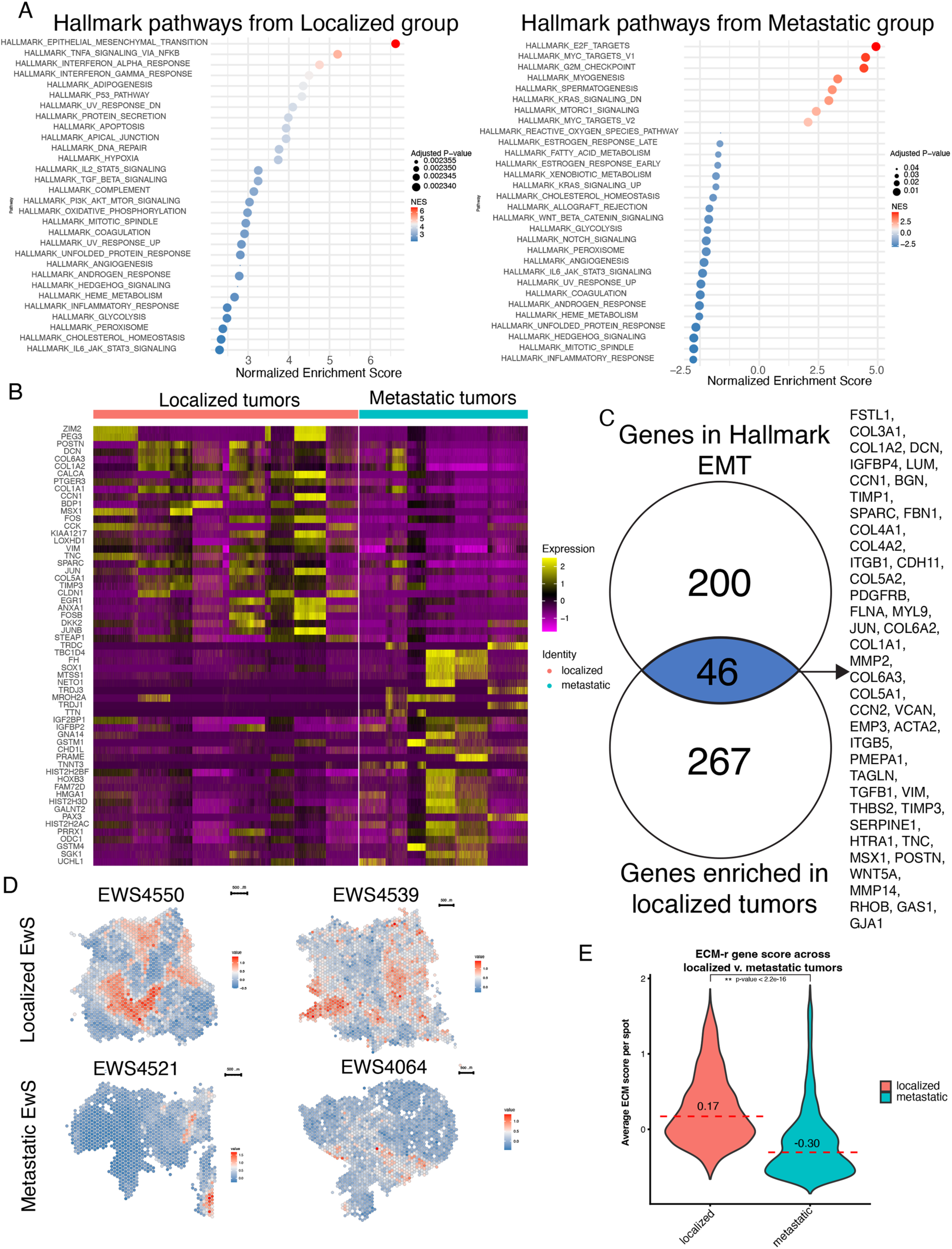
EwS tumors from patients with localized disease are enriched in extracellular matrix genes. (A) Gene set enrichment analysis (GSEA) for pseudo-bulk expression datasets of tumors from patients with localized and metastatic disease against Hallmark collection gene sets. (B) Heatmap of the most differentially expressed genes between pseudo-bulk expression EwS datasets from patients with localized and metastatic disease. (C) A set of 46 differentially expressed genes in tumors from patients with localized disease that overlap Hallmark epithelial mesenchymal transition gene sets. (D) Spatial mapping of spots enriched in the overlapping 46 extracellular matrix genes (ECM) on representative tumors (localized: EWS4550, EWS4539, and metastatic: EWS4064, EWS4521). (E) Distribution of the average calculated enrichment score per spot between tumors from patients with localized and metastatic disease.

To determine genes that are differentially expressed in loc-EwS and met-EwS, we performed FindAllMarkers (R Seurat) to assess for significantly differentially expressed genes within loc-EwS and met-EwS using Wilcoxon rank-sum test with an average log-fold change > 0.25 (Table 2). Loc-Ews tumors displayed higher expression of extracellular matrix (ECM) genes such as *DCN*, *COL6A3*, *COL1A2*, *COL1A*, *TNC*, *COL5A1*, *SPARC* (Figure 4B). In contrast, met-EwS, exhibited upregulation of genes associated with proliferation and metastasis, such as High-Mobility Group AT-Hook 1 (*HMGA1*), the cancer/testis antigen Preferentially Expressed Antigen in Melanoma (*PRAME)*, Glutathione S-Transferase Mu 1, and T cell receptors related genes. Previously, gene expression analysis from EwS patients that died identified several differentially expressed genes related to cell motility, cell migration and cell adhesion, and reported that *GSTM2*, a glutathione metabolism gene, was associated with worse outcome.^33^ We found that met-EwS tumors exhibited enrichment of genes associated with glutathione metabolism (*GSTM1, GSTM4, SMS* and *ODC1*) and cell proliferation (*TWIST1, IGFBP2, IGF2BP1*) (Table 2), along with higher expression of High-Mobility Group AT-Hook 1 (*HMGA1*) and the cancer/testis antigen *PRAME*. *HMGA1* was recently reported to be overexpressed and associated with trabectedin resistance in soft tissue sarcomas.^34,35^ *PRAME* has been proposed as a potential multi-pediatric cancer target, and noted to have robust protein expression in patient derived xenografts of EwS.^19^ Glutathione S-transferases (GSTs) are isoenzymes that have been implicated in developing chemoresistance in cancer, and have been noted to be a promising candidate to overcome drug resistance against conventional chemotherapy in osteosarcoma, EwS and rhabdomyosacoma.^36,37^

Based on the GSEA showing that loc-EwS tumors are enriched in genes in the EMT hallmark pathway, and the results from differential gene expression analysis indicating enrichment in ECM genes in loc-EwS, we next examined the overlap between the 267 differentially upregulated genes in loc-EwS and the 200 genes from the EMT hallmark pathway (Figure 4C). This analysis identified 46 overlapping genes, which we designated as ‘ECM-related’ (ECM-r) genes. To further explore their spatial distribution, we then performed AddModuleScore to calculate average expression levels of the ECM-r composite genes per spot across each ST tumor sample. Visualization of ECM-r gene enrichment revealed strong colocalization with stromal regions in both loc-EwS and met-EwS tumors (Figure 4D). However, loc-EwS had a higher ECM-r composite scores when compared to met-EwS using a Wilcoxon rank-sum test (Figure 4E). In summary, loc-EwS express ECM-r genes more highly compared to met-EwS and their spatial expression is concentrated within the tumor stroma, underscoring the distinct microenvironmental characteristics of localized EwS.

### Gene composite analysis demonstrates unique spatial localization of immune cells and ECM in EwS

Our finding that loc-EwS have more ECM-r genes compared to met-EwS, the complex relationship between the ECM and the immune system, and the spatial overlap between stroma and immune signatures, prompted us to look further into the spatial mapping of EwS tumor cells, stroma and immune cells.^38^ To map the spatial organization of these elements, we performed AddModuleScore analysis on EwS composite genes which consisted of *EWSR1::FLI1* target genes *NKX2-2*, *CD99, CAV1, CCND1, HES1, KDSR, PAPPA,*^16,39–41^ and found that the visualized spatial expression of this EwS composite score was consistent with the tumor distribution on H&E slides (Figure 5A,B, Supplementary Figure S2A-G). ECM-r genes had prominent expression in loc-EwS compared to met-EwS. As expected, ECM-r gene expression did not overlap with EwS composite genes.

**Figure 5:**
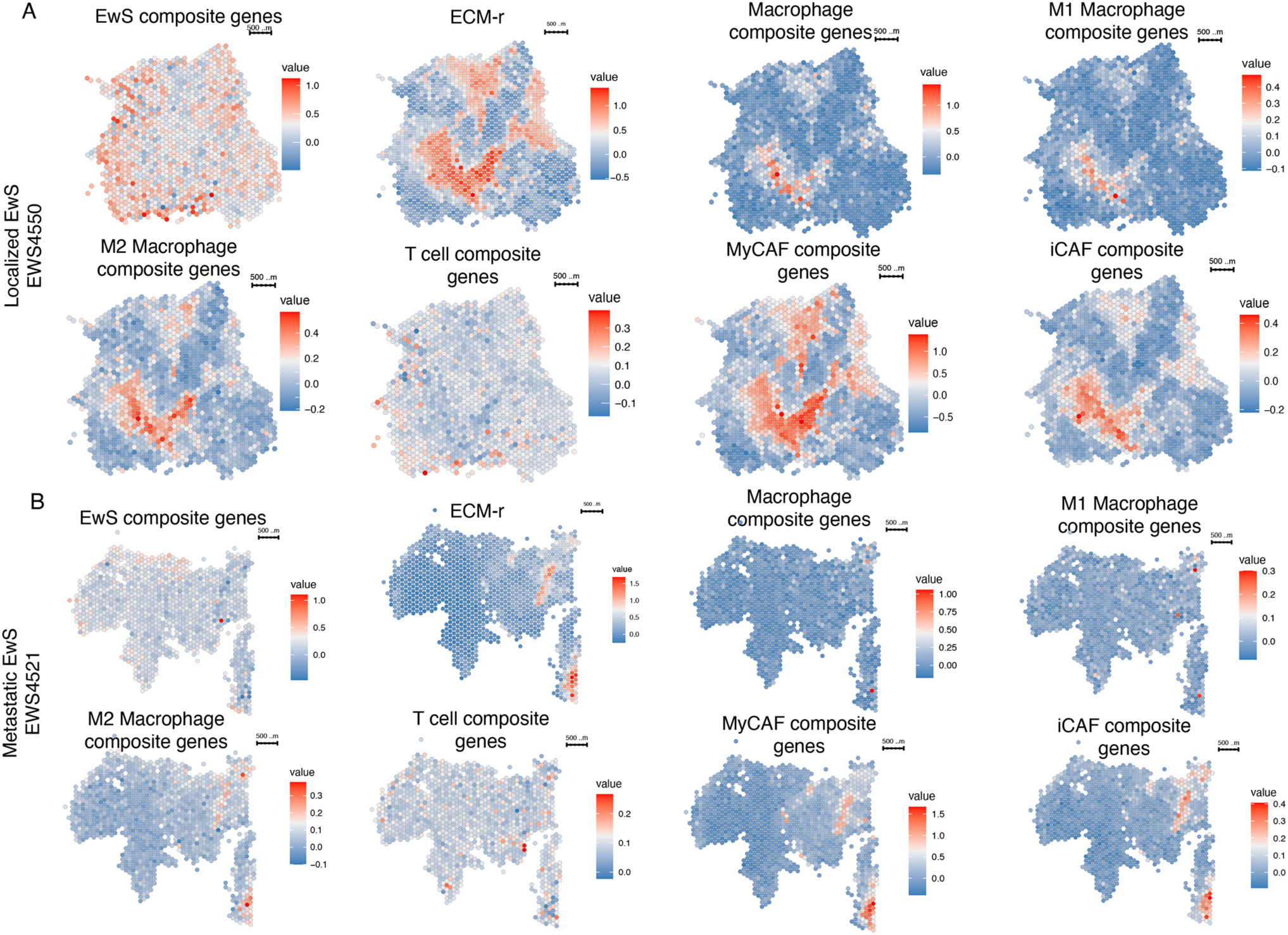
Unique spatial gene signatures reveal colocalization of macrophages with stromal regions of EwS tumors. (A) Representative spatial gene enrichment mapping of localized tumor EWS4550. Spatial mapping of spots enriched in composite genes that annotate EwS, ECM, and immune cells (macrophage, T cells, cancer-associated-fibroblasts) in EWS4550. (B) Representative spatial gene enrichment mapping of metastatic tumor EWS4521 with composite genes that annotate EwS, ECM, and immune cells in EWS4521.

Next, we assessed the presence of macrophage/monocytes using composite genes for macrophage/monocytes,^42^ M1 macrophages and M2 macrophages (Table 3).^43^ In both loc-EwS and met-EwS tumors, the macrophage gene composite colocalized with the ECM gene composite, though ECM gene expression was much more prominent in loc-EWS. Expression of T cell gene composite markers^44^ occurred predominantly within the EwS tumor spots compared to ECM-r spots (Figure 5 A,B). This spatial pattern was seen in both loc-Ews and met-EwS tumors. Due to the prominence of ECM-r regions, we assessed the presence of cancer-associated-fibroblasts (CAFs) specifically with myofibroblastic CAFs (myCAFs) and inflammatory CAFs (iCAFs) gene composites.^45^ Both myCAFs and iCAFs colocalized in regions enriched in ECM-r genes and regions enriched in M1 and M2 macrophages in both loc-EwS and met-EwS (Figure 5A,B) (Figure S3). Taken together, we demonstrated that spatial gene annotations of EwS, ECM and immune cells exhibit distinct spatial localization patterns within the tumors, however, with limited resolution to discern differences between loc-EwS and met-EwS.

### Spatial deconvolution reveals infiltrative T and B cells in Ewing sarcoma tumors

Recently Stahl and colleagues utilized CIBERSORT to deconvolute 197 microarray gene expression datasets of primary EwS tumor samples and reported that the most abundant cell types were predominantly immunosuppressive M2 macrophages, followed by T cells.^46,47^ Given these findings and the intriguing unique spatial localization of immune cells based on gene composite scores observed in our tumor cohort, we investigated immune infiltration in our cohort of EwS tumors, by performing CIBERSORTx.^24^ We generated pseudobulk RNA counts of each of the 16 tumors and performed CIBERSORTx.^24^ The predominant immune cell population were resting memory CD4 T cells (median: 27%), followed by M0 macrophage (median: 21%) and M2 macrophage (median:14%) (Figure 6A) (Table 4). However, CIBERSORTx profiled is restricted to 22 defined human immune cell types,^47^ thus potentially missing other relevant immune cell types. Additionally, CIBERSORTx does not provide spatial location of these immune cell types.

**Figure 6:**
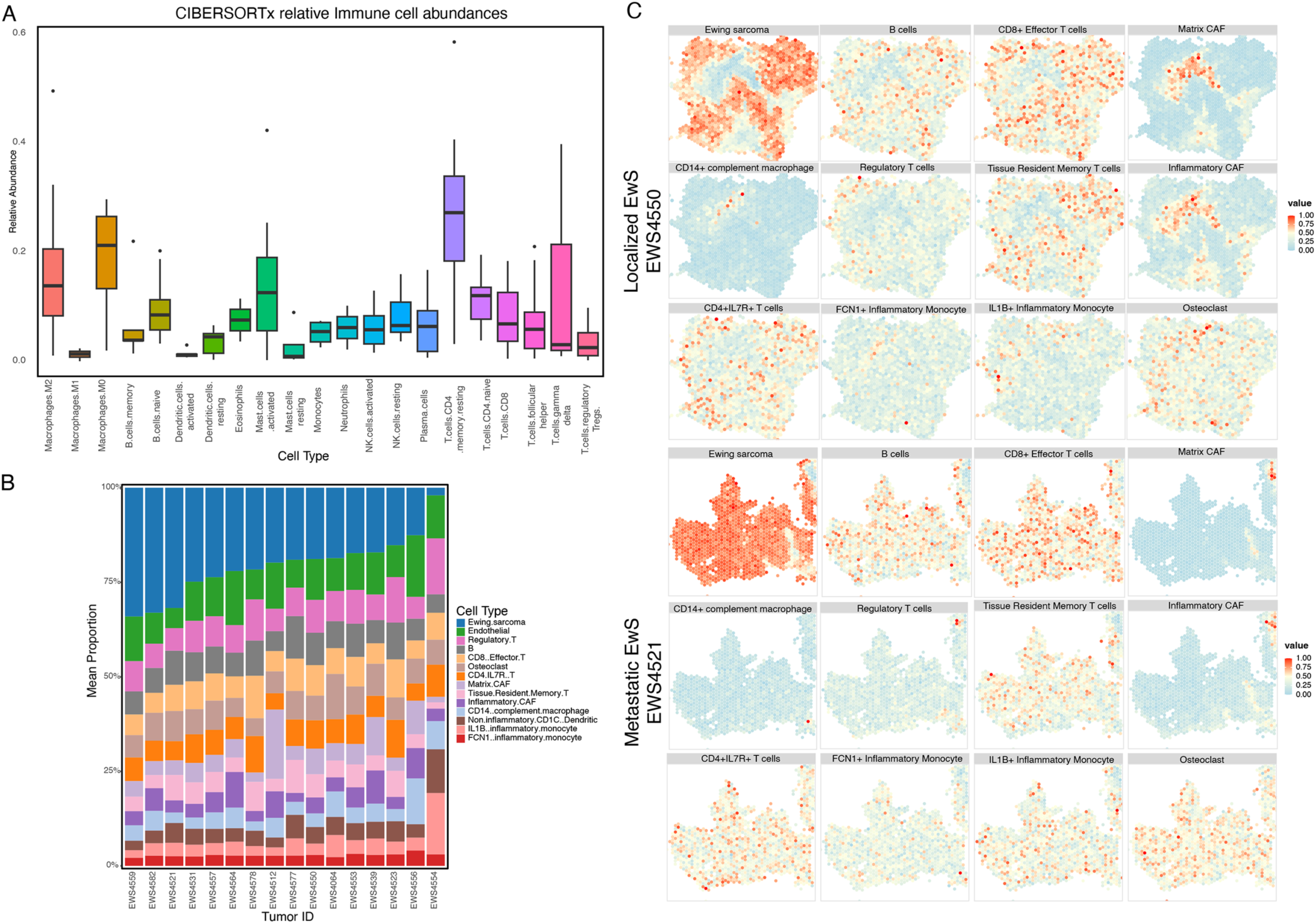
Deconvolution analysis demonstrates heterogeneous immune cell populations and distinct spatial localization of immune cells. (A) Estimation of immune cell abundances in all 16 tumor samples through CIBERTSORTx. (B) Deconvolution CARD analysis of all 16 tumor samples using a publicly available EwS scRNAseq dataset as a reference with individual boxplots. Each cell population is depicted in a different color. (C) Spatial visualization of deconvoluted populations from two representative tumors (EWS4550 (localized), and EWS4521 (metastatic)).

To understand the spatial distribution of the cell types within each of the 16 tumors, we leveraged publicly available scRNAseq datasets of EwS tumors and performed deconvolution analysis using CARD (Figure 6B).^23,22^ Deconvolution with the reference EwS scRNAseq dataset provided insight in the proportions of various cell types within each tumor (Figure 6B). Significant intertumoral heterogeneity is noted among all tumors, with each tumor having different proportions of immune cells. We then visualized the deconvoluted proportions spatially. As expected, the transcriptomic inference of EwS cells’ location recapitulated where EwS cells are localized within the H&E image in both loc-EwS and met-EwS tumors (Figure 6C, Supplementary Figure S3A-F). Similar to what we observed previously in gene composite analysis, we noted colocalization of CAFs at regions where ECM is dense, and T cells scattered throughout the tumor-rich regions (Figure 6C, Supplementary Figure S3A-F). Spatial deconvolution further revealed notable intratumoral heterogeneity within each immune cell subset. In addition to T cells, we identified previously undescribed infiltrative B cell populations, and various T cell subsets, including T regulatory cells (Tregs), CD8+ effector T cells, CD4+IL7R+ T cells and memory T cells. We also noted that monocytes and macrophages were mostly congregated near or in ECM dense regions, and colocalized with CAFs (Figure 6). Contrasting to the CD14+ complement macrophages, other myeloid cells such as IL1B+ inflammatory monocytes, FCN1+ inflammatory monocytes and osteoclasts were dispersed throughout the tumor. While enrichment of Tregs colocalized with enrichment of EwS tumor cells, the signal was not as prominent as CD8+ effector T cells. In summary, deconvolution of ST data enhanced the resolution of the EwS TME revealing previously unreported immune cell infiltrates, and enabled spatial mapping of their distribution. This analysis also reinforced the intratumoral and intertumoral heterogeneity of EwS tumors, providing deeper insights into the immune landscape of EwS.

### Loc-EwS tumors exhibit unique spatial ECM, and immune microenvironmental signals

Based on our previous findings that loc-EwS tumors are enriched in ECM related genes compared to met-EwS, and display unique spatial organization of immune cells through spatial deconvolution, we tested whether differences in TME contribute to biological differences between loc-EwS and met-EwS. To explore this, we first imputed the ST data using Adaptively-thresholded Low Rank Approximation (ALRA),^48^ and then performed NICHES on individual tumors, exploring the ligand-receptor (LR) signaling connectivity in the neighborhood of each Visium spot using LR lists from Omnipath.^25,49^ We integrated the NICHES results and clustered the data with a resolution of 0.2, resulting in five NICHES microenvironmental clusters (Figure 7A). These microenvironmental clusters exhibited a distinct pattern compared to the original tumors’ transcriptomic phenotypic clusters (Supplementary Figure S4). Microenvironmental cluster 0 primarily overlaid with the ECM regions of the tumor, while microenvironmental cluster 1 primarily corresponded to EwS dense regions. To further characterize these clusters, we performed FindAllMarkers and compiled a microenvironmental signaling list using only significant LR markers (p-value <0.001) (Table 5).

**Figure 7:**
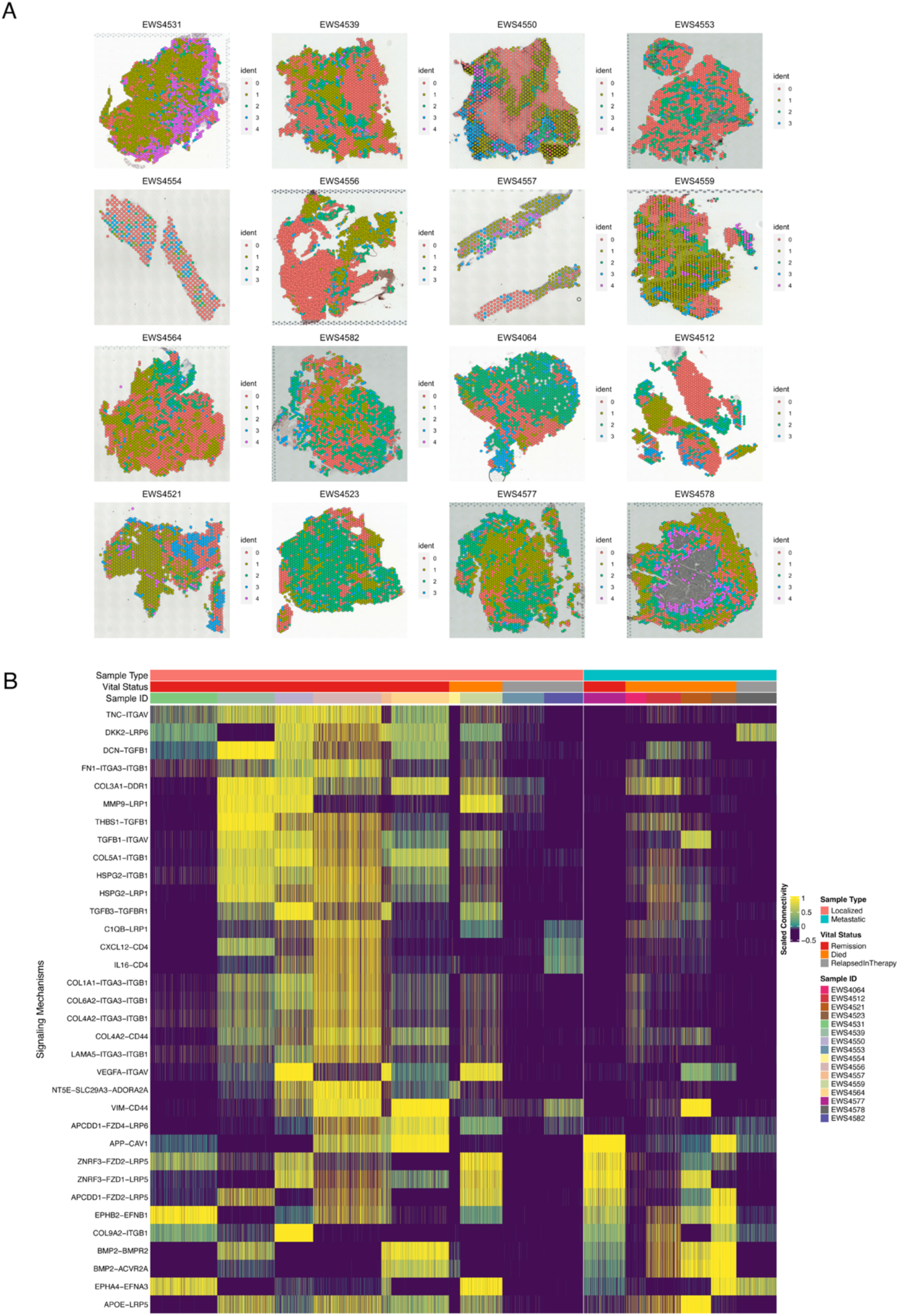
Spatial ligand-receptor microenvironmental transcriptomic signals in localized and metastatic EwS Tumors. (A). NICHES ligand-receptor neighborhood to spatial spot microenvironmental clustering identify 4 unique microenvironmental clusters. (B). NICHES analysis demonstrate enrichment of ECM-related microenvironmental connectivities in loc-EwS tumors.

Based on these observations, we specifically investigated the predominant microenvironment signaling connectivity unique to microenvironmental clusters 0 and 1, and examined the generated LR marker list (Table 5). NICHES analysis revealed unique microenvironmental connectivity signals in cluster 0 and cluster 1 (Supplementary Figure S5). In cluster 0, we observed significant ECM and immune related signals, such as collagen related LR microenvironmental signals (*COL4A1-CD93, COL4A1-ITB1, COL1A2-ITGB1)*, TGF-beta related signals (*TGFB1-SDC2, TGFB1-TGFBR1-TGFBR2),* immune-related signals *(CD14-ITGB1, CD14-RIPK1, CXCL12-CD4, IL16-CD4 and MIF-CD74)* (Supplementary Figure S5). In cluster 1, we observed predominant proliferative connectivity signals such as *WNT* related signals (*WNT5A-FZD1, WNT5A-LRP5, APCDD1-FZD1-LRP5)*, Fibroblast Growth Factor Receptor (FGFR) signaling *(FGF13-FGFR4*), and immune signals *(CTF1-LIFR*) (Supplementary Figure S5).

Our finding that microenvironmental clusters hold unique microenvironmental connectivity signals led us to examine further what distinct signals exist within loc-EwS tumors compared to met-EwS tumors. We generated a loc-EwS LR marker list (Table 6) and a met-EwS marker list (Table 7). Loc-EwS tumors had prominent ECM related microenvironmental signals compared to met-EwS, such as collagen-related interactions and heparan sulfate proteoglycan-related interactions (Figure 7B). We also noted that *TGFB* signals were enriched in loc-EwS compared to met-EwS. We observed *WNT* signaling activation in both loc-EwS and met-EwS tumors, with an enrichment of microenvironmental connectivity associated with *WNT* inhibition signaling pathways in both loc-EwS and met-EwS tumors, such as *APCDD1-FZD4-LRP6* and *APCDD1-FZD2-LRP5.* Loc-EwS predominantly exhibited immune signaling pathways such as *C1QB-LRP1, CXCL12-CD4, and IL16-CD4*, in contrast to met-EwS (Figure 7B). In summary, loc-EwS tumors are enriched in ECM related microenvironmental signals in contrast to met-EwS tumors, and loc-EwS tumors are enriched in immune signaling compared to met-EwS tumors. These differences underscore the distinct microenvironmental landscapes of localized and metastatic EwS, highlighting potential targets for therapeutic intervention.

### Ultrahigh-plex spatial phenotyping reveals B cells, lymphoid aggregates, and infiltrative M2 macrophages in EwS tumors

Given that the immune signal *MIF-CD74* was noted to be expressed in all tumors and may play a role in immunosuppression within the EwS TME, we investigated this signal further. To validate the unique spatial organization of immune cells, spatial enrichment of ECM and immune-related microenvironmental signaling of *MIF-CD74* reported in ST, we performed ultrahigh-plex PhenoCycler analysis on available EwS samples.^26^ Sufficient tissue was available for spatial proteomics in 14 of the 16 EwS tumors on which we previously performed ST. We confirmed our deconvolution results and localized the infiltrative B cell populations in both loc-EwS and met-EwS tumors (Figure 8A,B). CD8+ T cells were detected in all tumors, with higher density in loc-EwS tumors compared to met-EwS tumors (Figure 8A and Supplementary Figures S6A,C,E,G,I,K). Consistent with prior reports, the predominant leukocyte population in the TME were tumor-associated macrophages (TAMs) (Figure 8A,B).^46^ The majority of the TAMs surrounding CD8+ T cells expressed M2 markers (CD163+ and CD206+) rather than M1 (CD68+),^50,51^ and direct cell-cell contacts between M2 TAMs and CD8+ T cells were visualized in all EwS tumors (Supplementary Figures S6A,C,E,G,I,K). Furthermore, we discovered dense aggregates of T- and B- cells with TAMs in unique spatial locations within both loc-EwS and met-EwS tumors (Supplementary Figure S7).

**Figure 8:**
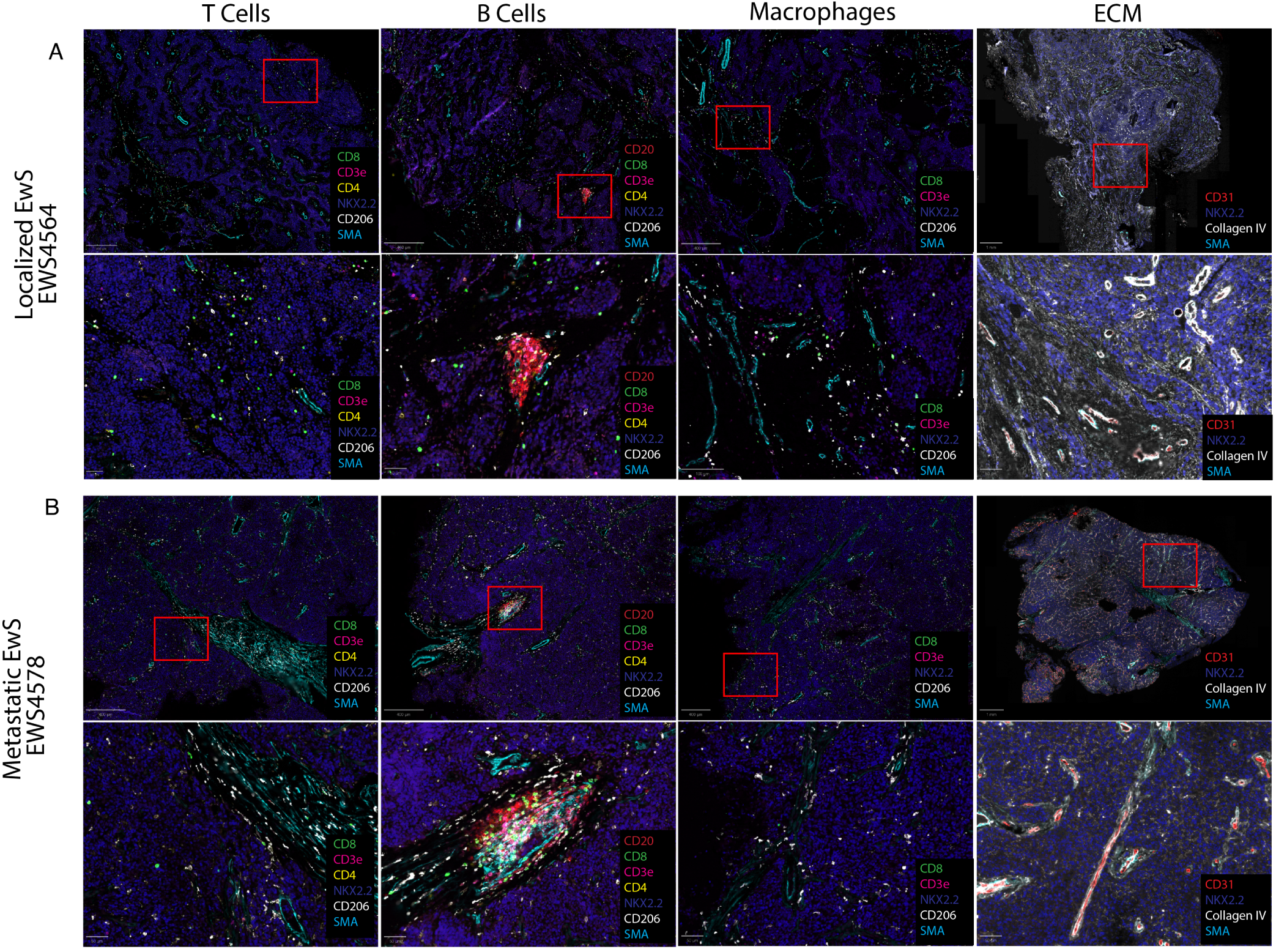
Ultrahigh-plex single cell spatial proteomic analysis of immune cells in EwS TME. (A) Single-cell spatial proteomic analysis demonstrates presence of T-cells, B-cells, macrophages and enrichment of ECM in a representative Loc-EwS tumor (EWS4564). (B) Single-cell spatial proteomic analysis demonstrates presence of T-cells, B-cells, macrophages and enrichment of ECM in a representative Met-EwS tumor (EWS4578).

TAMs are infiltrative in solid tumors, and are highly dynamic and heterogeneous.^52^ TAMs have the ability to polarize to a pro-inflammatory (M1) state or a pro-tumoral (M2) state in response to environmental perturbations.^52^ We observed an abundance of infiltrative M2 TAMs (CD163+ and CD206+) in EwS tumors compared to M1 TAMs (CD68+). Consistent with our ST results, in addition to being infiltrative in tumors, TAMs co-localized with tumor-associated stroma (collagen IV) (Supplementary Figure S6B,D,F,H,J,L). Loc-EwS tumors exhibited greater stromal enrichment (Collagen IV) compared to met-EwS tumors (Figure 8A,B). Additionally, we confirmed the protein expression of TGF-B1 in both loc-EwS and met-EwS. Finally, we validated the immune-related signals PD1-PD-L1 and MIF-CD74 (Figure 9A,B). Based on the abundance of M2 TAMs, we investigated the phenotype of TAMs within the environment of tumor-associated MIF. MIF was highly expressed in all EwS tumors, with M2 TAMs expressing CD74 clustering in areas of high MIF microenvironments (Supplementary figure S6B,D,F,H,J,L). Interestingly, spatial regions with less tumor-associated MIF expression harbored more M1 TAMs (Supplementary figure S6B,D,F,H,J,L). Taken together, these results suggest that MIF-CD74 signaling between EwS and TAMs may drive or potentiate the immunosuppressive M2 predominant TME in EwS tumors.

**Figure 9:**
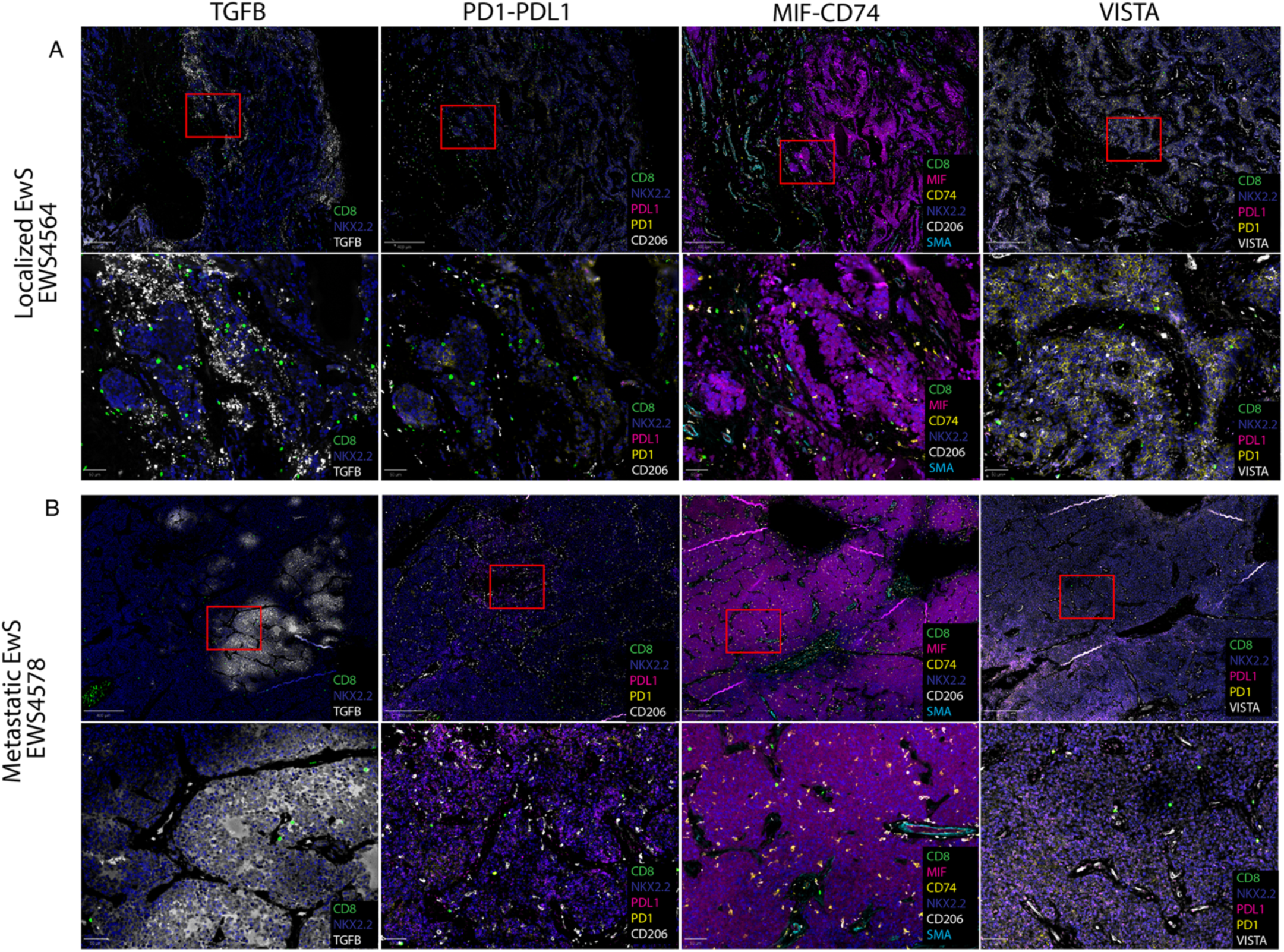
Ultrahigh-plex single cell spatial proteomic analysis of immune microenvironmental signals in EwS TME. (A) Single-cell resolution of TME modulating signals with TGFB1, PD1-PDL1, MIF-CD74 and VISTA in a representative Loc-EwS tumor (EWS4564). (B) Single-cell resolution of TME modulating signals with TGFB1, PD1-PDL1, MIF-CD74 and VISTA in representative Met-EwS tumor (EWS4578).

## Discussion

The ECM is an integral component of the TME that is highly diverse, acting as a reservoir for pro-tumorigenic and anti-tumorigenic cytokines, serving as communication network between surrounding cells, and functioning as an anchor for tumor migration/invasion and as a mechanical barrier against tumor growth.^53–56^ Previous efforts to identify prognostic gene profiles in tumors from patients with localized EwS highlighted variation in stromal content, with stroma-rich tumors expressing integrin and chemokine gene sets linked to survival.^33^ While tumor stroma was traditionally viewed as an immune barrier, it is now increasingly redefined as an immune rheostat, exerting context specific pro- and anti-tumor effects.^57^

In this study, we discovered differentially expressed ECM-r genes in loc-EwS tumors that were spatially enriched primarily in the stromal regions including several collagen genes (*COL1A2*, *COL3A1*, *COL4A1*, *COL5A2*, *COL6A3*), matrisome genes (*SPARC, TNC, FBN1*), and immune genes (*HLA-A, HLA-E*). Individually these ECM-r genes were expressed at unique spatial locations of the tumors, suggesting site-specific tumor-immune-stroma interface. To examine potential effects of the ECM on immune cell infiltration in EwS tumors, we assessed the spatial distribution of tumor and immune cell genes, and enhanced the resolution through deconvolution, employing reference EwS scRNAseq data.^23^ Matrix and inflammatory CAFs colocalized with macrophages, specifically CD14+ complement macrophages at the stromal region. Multiple T cell subsets were enriched where EwS tumors were enriched as well, such as CD8+ T cells and tissue resident memory T cells, juxtaposing the stromal region. Additionally, there was a notable presence of B cells inferred in spots coinciding with EwS tumor cells, a previously undescribed finding in the TME of EwS. Across all 16 tumors, B cells and CD8+ T cells did not colocalize with CAFs. Interestingly, majority of the loc-EwS tumors (60%) had spot correlation of cytotoxic CD8+ T cells with EwS cells, in contrast to met-EwS tumors (33%) (Supplementary figure S8A-C), further reinforcing the unique tumor immune microenvironment (TIME) in the ECM enriched loc-EwS tumors.

To study the TME signaling connectivities underlying the stromal and tumor regions between loc-EwS and met-EwS tumors, we performed NICHES analysis. We discovered that loc-EwS tumors harbored unique spatial ECM, immune and proliferative microenvironmental signals. A key finding was the enrichment of TGF-β, a well-established potent regulator of the TME.^58,59^ *TGFB1-TGFBR1-TGFBR2* and *TGFB3-TGFBR1-TGFBR2* interactions were noted to be expressed in majority of the loc-EwS predominantly enriched in the stromal regions (Supplementary figure S9). Furthermore, we noted *TGFB1-SDC2* microenvironmental signal in majority of the loc-EwS tumors. Functionally, SDC2 acts as a co-receptor for TGFBR to mediate TGFB signaling, increasing ECM deposition and fibrosis, aligned with our ECM-enriched loc-EwS tumors.^60,61^ Several collagens-integrins interactions were also noted within ECM-enriched cluster, including *COL4A1-ITGB1, COL1A2-ITGB1, COL6A1-ITGB1*, suggestive of integrin related immune cell adhesion and immune response.^62,63^ Di Martino et al recently reported that tumor derived collagen III interacts with DDR1 (discoidin domain receptor tyrosine kinase) to maintain a wavy collagen architecture, thereby restricting cancer cell proliferation in a breast cancer model.^64^ Interestingly, we discovered significant enrichment of COL3A1-DDR1 connectivity predominantly in loc-EwS tumors, suggestive of stroma restricting tumor growth. In loc-EwS tumors, we noted significant enrichment of immune cell recruitment signals (CXCL12-CD4 and IL16-CD4) within the ECM enriched cluster. CXCL12 is known to play a critical role in monocyte extravasation, macrophage differentiation and can stimulate antigen-specific T cell responses.^65^ IL16 is a chemoattractant for CD4-immune cells such as T cells and monocytes/macrophages, and induces production of inflammatory cytokines on monocytes and antigen presenting cells.^66–69^ We also noted signals of macrophage migration (HSPG2-LRP1), and proinflammatory activation (CD14-RIPK1) within the ECM enriched regions.^70,71^ These findings highlight the anti-tumorigenic role of stroma and its involvement in immune cell recruitment and activation within the stroma of EwS tumors.

Despite evidence of immune cell recruitment and proinflammatory activation in our tumors, the TME of EwS is typically classified as immune-cold, with generally poor responses to immunotherapy.^72^ This raises the question of whether other mechanisms within EwS are counteracting the immune activation signals within the stromal compartment, thereby rendering immune surveillance dysfunctional. To investigate this further, we observed MIF-CD74 immune signaling expressed in all tumors. MIF is a proinflammatory cytokine that is overexpressed in several cancers such as lung cancer, osteosarcoma, neuroblastoma and melanoma.^73^ MIF is known to bind to CD74 with a role in immune homeostasis, cancer proliferation and inhibition of apoptosis.^74^ Importantly, MIF is implicated in immune evasion, driving the polarization of M2 macrophages and inhibiting T cell activation.^75–77^ Additionally we also found an IL1RAP-PTPRF signaling pathway that was expressed in all tumors. IL1RAP is known to be induced by EwS and suppresses anoikis to metastasize.^78^ These spatially unique immune related TIME signals highlight the complex and novel mechanisms that EwS may utilize to manipulate its TME, enabling tumor proliferation and immune evasion. Additionally, we noted PDL1-PD1 was only expressed in <25% of tumors, with signals predominantly in loc-EwS tumors. This may explain why, to date, PD-1 inhibitor treatment has not yielded clinical responses in Ewing sarcoma.^72^

There is a dearth of literature investigating in the expression of CD8+ T cells in EwS tumors. A study by Machado and colleagues reported the lack of infiltrative CD8+ T cells in EwS tumors.^79^ Consistent with this, our ultrahigh-plex spatial proteomic analysis revealed a scarcity of CD8+ T cells within the TIME of met-EwS compared to loc-EwS. Interestingly, EwS tumors had a striking abundance of M2 TAMs infiltrating both the stromal and tumor compartments in direct contact with the surrounding CD8+ T cells and EwS cells. Despite M2 TAMs being the predominant TAM subtype in EwS tumors, the detailed functional role of TAMs within the EwS TME remains poorly understood. In pancreatic ductal adenocarcinoma, TAMs exhibited characteristics of both M1 and M2 macrophages, holding both anti- and proinflammatory properties.^80^ Thus, the M2 TAMs within the TME of EwS may still retain antigen presenting capabilities. In recent years, a deeper understanding of lymphoid aggregates and tertiary lymphoid structures (TLSs) have improved the response to immunotherapy in lung cancer.^81,82^ Currently, there are no reports of lymphoid aggregates in EwS tumors. The presence of dense lymphoid aggregates in our EwS tumors raises the question of the possible function and the prognostic association of lymphoid aggregates in the TIME of EwS.

This is the largest study to date of ST analysis on treatment naïve EwS tumors obtained at diagnosis composed of both loc-EwS and met-EwS tumors. Previous efforts in profiling the immune composition of EwS have been limited based on microarray gene expression datasets, multiplex IHC, and scRNA-seq.^17,46^ While this study provides valuable insights into the EwS TME, we acknowledge some limitations. First, spatial transcriptomics performed via Visium V1 (10X) lacks a single cell resolution which may obscure finer cellular interactions. However, we attempted to mitigate this challenge via use of high-resolution multiplex immunostaining. Additionally, we did not have matched scRNAseq data from the same patients thus limiting our ability to perform direct transcriptional deconvolution. We primarily relied on clinical biopsy samples, which may not fully capture the heterogeneity of entire tumors. The small sample size of our cohort limited the statistical power to identify significant correlations between ST data and clinicopathological variables such as disease status/clinical outcome, age, or gender. We also acknowledge the need for external validation in a larger independent patient cohort to address the sample size limitation.

In conclusion, consistent with other studies, we found that EwS tumors exhibit significant intra- and inter-tumoral spatial heterogeneity.^16,17,23,83^ Our data revealed variable stromal representation, with greater stromal enrichment in loc-EwS compared to met-EwS. Additionally, we observed a scarcity of cytotoxic T cells in tumor compartment of met-EwS compared to loc-EwS. This suggests that the stromal component of EwS tumors may play an anti-tumorigenic role by acting as an immune recruitment center. The stromal compartment of EwS appears to harbor both innate and adaptive immune cells, promoting antigen presentation to re-direct T cells into the tumor. The reduced stromal component in met-EwS tumors results in the loss of this antigen presentation niche, specifically mediated by macrophages, thereby contributing to immune escape. Furthermore, our data also emphasize the multidimensional complexity of the TIME in EwS. EwS tumors employ various immune signals to evade immune surveillance and convert the surrounding TME into an pro-tumorigenic state, such as through MIF-CD74, and VISTA. These signals occur simultaneously in various spatial regions of EwS tumors, which may explain the limited success of mono-immunotherapy approach thus far.

Additionally, the presence of CD8+ T cells within an M2 TAMs predominant TIME raises the question of whether EwS has a hot TME instead, with the CD8+ T cells’ presence being neglected due to the complex immune evading mechanisms in the TME.^84^ Most importantly, we report previously undescribed novel immune targets that may revert the pro-tumor TME to an anti-tumor TME. Future studies incorporating whole-tumor mapping and higher-resolution transcriptomic methods will be essential to expand on these findings. Further investigation of the immunomodulatory role of stroma in EwS as well as the mechanistic roles of MIF-CD74 signaling within EwS TME in vitro and in vivo may uncover novel therapeutic opportunities to counteract immune evasion and improve EwS patient outcomes.

## Methods

### Sample selection

A total of 16 formalin-fixed paraffin-embedded treatment naïve primary Ewing sarcoma tissue samples were obtained from patients with localized or metastatic Ewing sarcoma at diagnosis. Samples underwent quality check process, and all samples had DV200 scores >/= 30-50% prior to being selected for spatial transcriptomics (Table S1). The clinicopathologic characteristics of samples used in the study were summarized in Table 1. All samples and clinical data were obtained from the Children’s Hospital Los Angeles (CHLA) Tumor Registry Study under protocols approved by the CHLA’s Institutional Review Board (CCI-06-00198) Written informed parental permission, and where possible, child assent, was obtained for all patients participating in the study.

### Visium spatial transcriptomics

Spatial transcriptomics was performed according to the manufacturer’s protocol (10x Genomics). In brief, a 5-μm-thick EwS tissue FFPE sections were placed onto a spatial transcriptomics expression slide (10x Genomics, #1000188). Each gene expression slide contains four 6.5 mm x 6.5 mm capture areas covering 4992 spatially barcoded spots (resolution 1-10 cells/spot). Tissue sections on Visium slides were first fixed in methanol (Millipore Sigma #34860) followed by an aqueous eosin-based hematoxylin and eosin (H&E) staining, and imaging protocol CG000160). Brightfield images were acquired with a DMI6000B microscope equipped with a 20x/0.7 HC PL APO lens and a DFC295 color camera (Leica Microsystems, Buffalo Grove, IL). Following imaging of H&E staining, permeabilization, cDNA synthesis and library generation was performed according to manufacturer’s protocol for the Visium Spatial Gene Expression Slide and Reagent Kit (10x Genomics). Quality control of dual-indexed barcoded libraries was performed with Agilent Bioanalyzer. Paired-end dual indexed final libraries were sequenced with 1% Phix on the Novaseq platform with these parameters: Read 1-28 cycles, Index 1-10 cycles, Index 2-10 cycles, Read 2-90 cycles.

### ST data preprocessing and normalization

Sequencing ST data were preprocessed using the Space Ranger pipelines (version 1.3.1; 10x Genomics). Base call files (BCL) data were demultiplexed and converted to FASTQ files. The spaceranger count pipeline was then used for read alignment to human reference genome GRCh38, unique molecular identifier (UMI) counting, and generation of feature-spot matrices corresponding to the H&E image within the fiducial frame of the gene expression slide. The raw UMI counts and spatial barcodes were imported to R program STUtility (version 1.1.1).^20^ Gene expression count data were then normalized using the NormalizeData function from the R package Seurat (version 5.1.0).^21^

### Histopathologic annotation

FFPE tissues from each block selected for study were stained with H&E and non-necrotic tumor dense regions were manually selected by an expert pathologist for spatial transcriptomics. Images captured from Visium slides were reviewed and annotated by an expert pathologist.

### ESTIMATE analysis

ESTIMATE analysis was performed with the R package tidyestimate (version 1.1.1).^27^

### Gene set enrichment analysis

Gene set enrichment analysis (GSEA) was applied at the merged processed objects of all 16 tumors using the R packages msigdbr (version 7.5.1) and fgsea (1.30.0). We chose to perform GSEA on the merged object instead of integrated object because too much biological variation was removed upon integration process.

### Gene composite score annotation

Gene signatures were compiled to annotate EwS, Macrophages, M1 Macrophages, M2 Macrophages, T Cells, myCAFs and iCAFs (Table 2). The gene signatures were then used to generate a gene composite score per spot using the function AddModuleScore in Seurat (version 5.1.0).^21^

### CIBERSORTx analysis

Immune cell abundance estimation was performed through CIBERTSORTx.^24^ Normalized gene expression data was compiled to create a pseudobulk profile. This profile was then compiled as a mixture file and analyzed against the LM22 immune.^47^

### CARD Deconvolution analysis

Processed and normalized gene expression count data objects from STUtility were deconvolute through CARD using reference EwS scRNAseq dataset.^20,22,23^

### Ligand-receptor interaction analysis

Processed and normalized gene expression count data objects from STUtiity from all 16 tumors were merged, integrated and imputed with ALRA.^48^ Spatial ligand-receptor neighborhood to spot connectivity was assessed with NICHES on R using LR from Omnipath.^25,49^ Integrated NICHES results were then clustered with a resolution of 0.2 and predominant microenvironmental clusters 0 and 1 were selected for further downstream analysis and ligand-receptor (LR) marker list generation. LR marker list was generated based on microenvironmental clusters 0 and 1 and thresholds of average pct1 >0.8, average metastatic score >5, and average localized score >8 were selected in order to capture highly significant LRs that is present in majority of the tumors. To find significant mechanisms differentiating loc-/met Ews samples, we ran a statistical test for each possible 2-sample pairing, one sample from localized samples and the other from metastatic samples, with total of 10×6 = 60 pairs. In each statistical test we used FindAllMarkers() function in Seurat package, setting min.pct = 0.05, logfc.threshold=0.1 and only.post=T. Then we concatenated the 60 marker lists and counted the frequency of LR mechanisms, defining this frequency as score for that mechanism. Similarly, we define number of times the mechanism appeared as a Localized marker as “avg_Localized_score”, while number of times the mechanism appeared as a Metastatic marker as “avg_Metastatic_score”. During the aggregation of this concatenated list, we also generated new p-values and logFC values. We use Fisher’s method to combine old p-values. New logFC values are calculated by log_2_(mean(2^abg_log_2_FC)). An additional ratio score was generated by dividing avg pct1/avg pct2 score and a power score was generated by avg pct1/avg pct2*avg_log_2_FC to further assess most expressed LR pairs. Additionally, a loc-EwS LR marker list and a met-EwS marker list were generated with thresholds of average pct1 >0.8, average score >35 were selected for loc-EwS marker list, and thresholds of average pct1 >0.8, and average score >30 were selected for met-EwS marker list.

### Phenocycler-Fusion FFPE staining

FFPE sections of EwS tumors (5-μm-thick) were stained and barcode-conjugated antibodies (Table S2) according to the protocol offered by AKOYA biosciences (Phenolmager Fusion user guide PD-000011 REV M, Date: 12.05.2023) with minor modifications. In brief, sections were baked at 60°C overnight, dewaxed by incubating for 10 min in xylene (Sigma-Aldrich, 185566-1L) twice, hydrated in solutions of decreasing EtOH concentration, and washed for 5 min in 18.2MΩ water twice. They were then incubated in a NxGen Decloaking chamber (Biocare Medical DC2012) with 1x antigen retrieval AR9 solution (AKOYA, AR9001KT) for 20 min at 110°C. Slides were then allowed to sit at RT for 30 minutes and were then washed with 18.2MΩ water, followed by two washes in hydration buffer (AKOYA staining kit 7000017). Samples were then incubated in staining buffer (AKOYA staining kit 7000017) for 30 min, followed by staining with the antibody cocktail for 3 hours at RT. Stained slides were then washed in staining buffer, fixed with 1.6% Paraformaldehyde (EMS 15710 ) prepared in storage buffer (AKOYA staining kit 7000017) for 10 min at RT, washed 3x in PBS, treated with ice-cold methanol (Sigma-Aldrich 34860-1L-R) for 5 min at 4°C, washed 3x in PBS, fixed with fixative reagent (AKOYA staining kit 7000017) for 20 min at RT, washed 3x in PBS and stored in storage buffer at 4°C. Within 5 days of staining, antibody-stained slides were imaged using the reciprocal fluorescently labelled reporters (Table S2) on a Phenocycler-Fusion 2.0 system.^26^ Output images were segmented using the Steinbock pipeline.^85^

## Supporting information

Supplementary Images

## Acknowledgements

We thank the Children’s Hospital Los Angeles Pathology (Julie Galindo, William Gordo, and Monica Mendez), Spatial Biology and Genomics (Long Hung, Michael Hadjidaniel, Xiangming Ding, and Shahab Asgharzadeh), and Cellular Imaging (G. Esteban Fernandez) Cores. CK is supported by the St. Baldricks Foundation as the Shohet Family Fund St. Baldrick’s Fellow, the A.P. Giannini Foundation, and the Coco Johnson Fellowship in Pediatric Sarcoma. JFA is supported by grants from the National Cancer Institute (U54CA268072 and P30CA014089) and by the Alfred E. Mann Family Foundation Chair in Cancer Research. MSBR is supported by grant T32GM086287 from the National Institute of General Medical Sciences (NIGMS), and startup funds from the Yale School of Medicine. This study was partly supported by the National Institutes of Health P30CA046934 by utilizing the Bioinformatics and Biostatistics Shared Resource. The opinions expressed are those of the authors and do not necessarily represent the thoughts or opinions of NIGMS, NIH, or the United States government. This work is dedicated to Noah Shohet, in loving memory of Noah Shohet and Stanley Kuo.

## Author contributions

Christopher Kuo, Conceptualization, Funding, Data curation, Formal analysis, Investigation, Methodology, Validation, Visualization, Writing – original draft, Writing – review and editing; Krinio Giannikou, Formal analysis, Writing – Review and editing; Nuoya Wang, Formal analysis, Investigation, Methodology, Validation, Writing – review and editing; Mikako Warren, Formal analysis, Investigation, Validation, Writing – review and editing; Andrew Goodspeed, Data curation, Writing – review and editing; Nick Shillingford, Formal analysis, Writing – review and editing; Masanori Hayashi, Data curation, Writing – review and editing; Micha Sam Brickman Raredon, Formal analysis, Investigation, Methodology, Validation, Writing – review and editing; James F Amatruda, Conceptualization, Funding, Investigation, Methodology, Project administration, Supervision, Validation, Writing – review and editing.

