## Supplementary Images for "Tumor-associated stroma shapes the spatial tumor immune microenvironment of primary Ewing sarcomas"

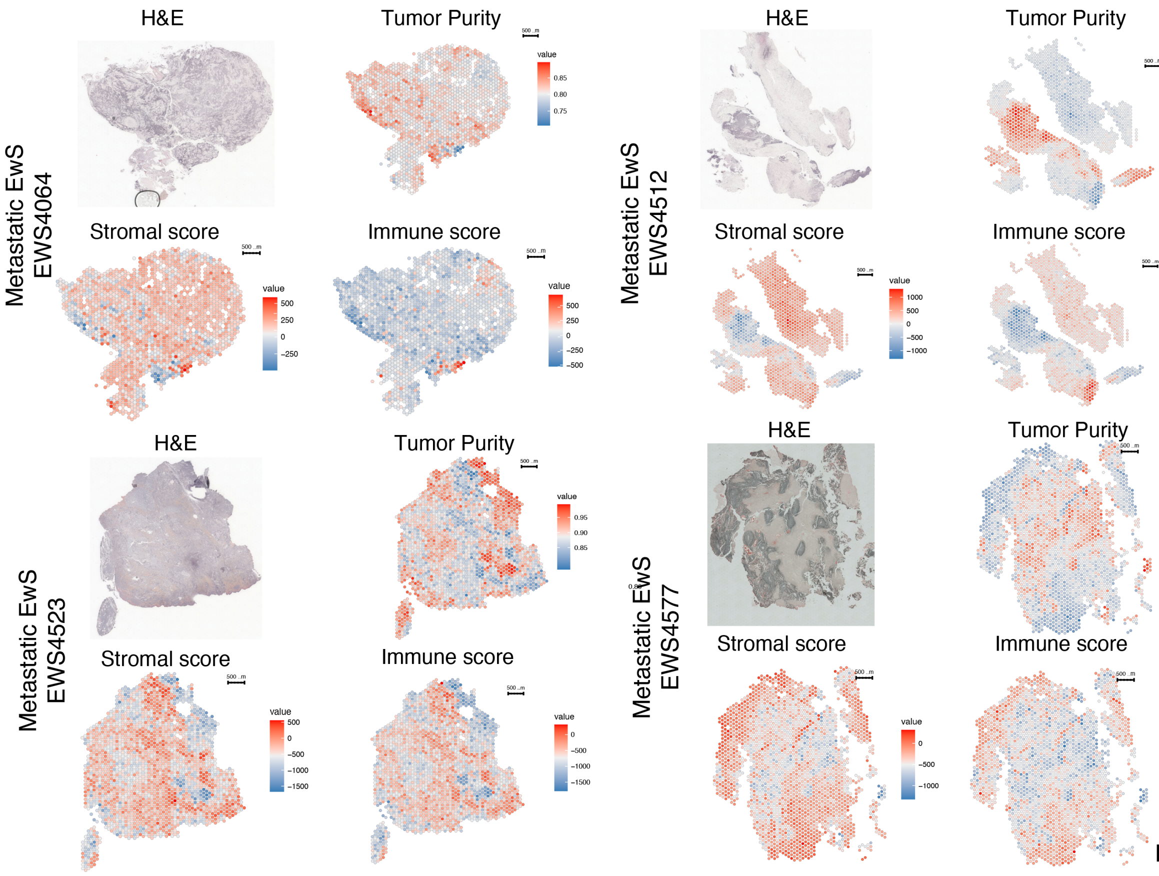

Metastatic EwS
EWS4064
H&E
Tumor Purity
Stromal score
Immune score
Metastatic EwS
EWS4512
H&E
Tumor Purity
Stromal score
Immune score
Metastatic EwS
EWS4523
H&E
Tumor Purity
Stromal score
Immune score
Metastatic EwS
EWS4577
H&E
Tumor Purity
Stromal score
Immune score
value


Fig.S1A

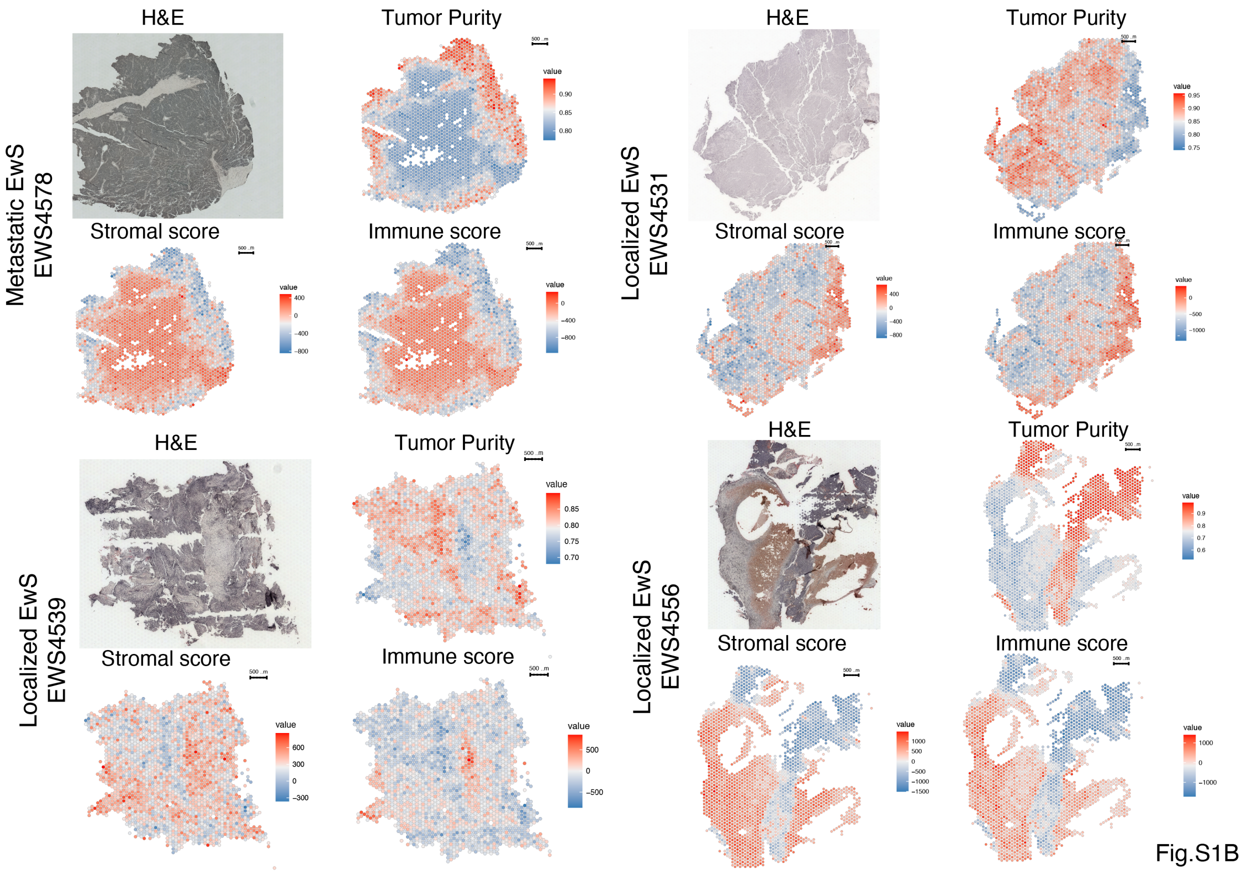
H&E
Tumor Purity
Stromal score
Immune score
Metastatic EwS EWS4578
Localized EwS EWS4531
Localized EwS EWS4539
Localized EwS EWS4556
value
0.90
0.85
0.80
400
0
−400
−800
0.95
0.75
−500
−1000
0.70
600
300
−300
500
0.9
0.8
0.7
0.6
1000
−1500

Fig.S1B

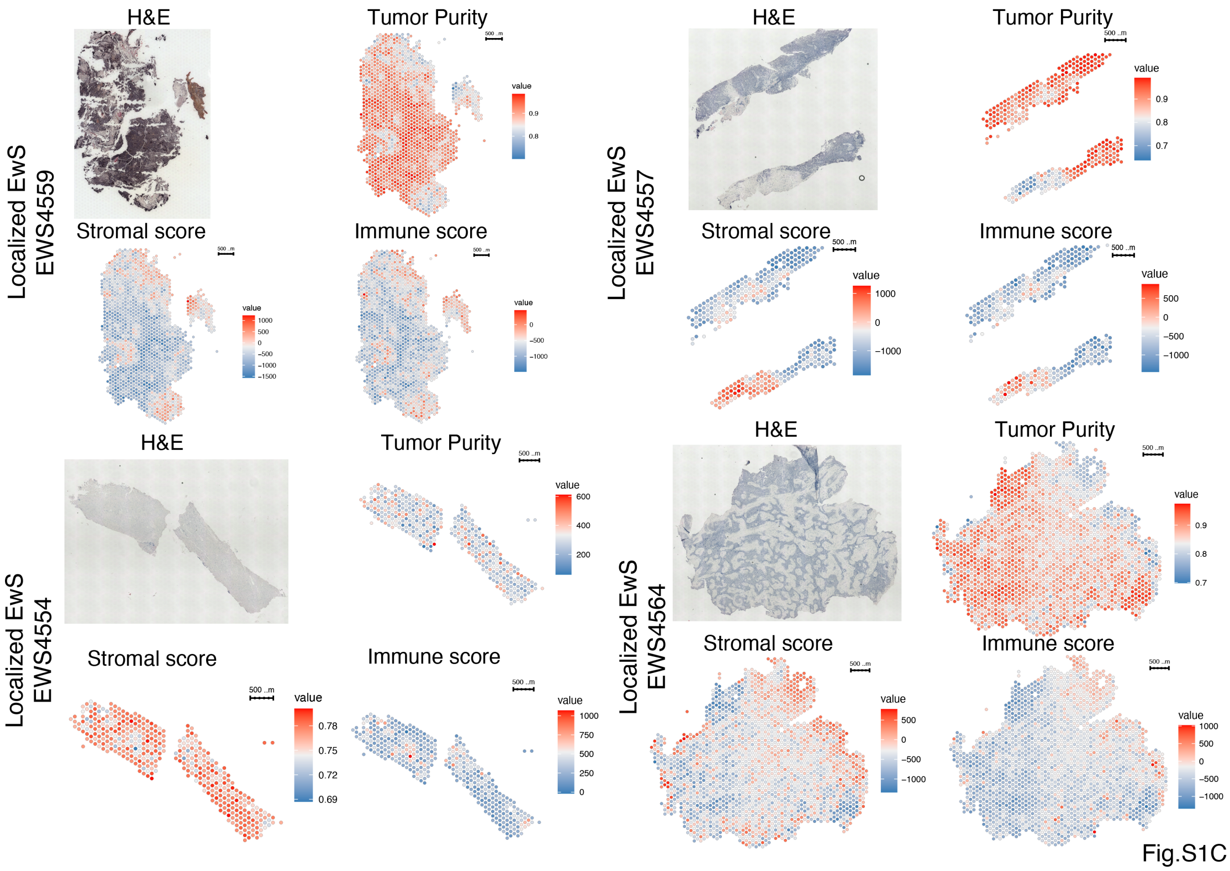
Localized EwS EWS4559
H&E
Tumor Purity
Stromal score
Immune score
Localized EwS EWS4557
H&E
Tumor Purity
Stromal score
Immune score
Localized EwS EWS4554
H&E
Tumor Purity
Stromal score
Immune score
Localized EwS EWS4564
H&E
Tumor Purity
Stromal score
Immune score
value

Fig.S1C

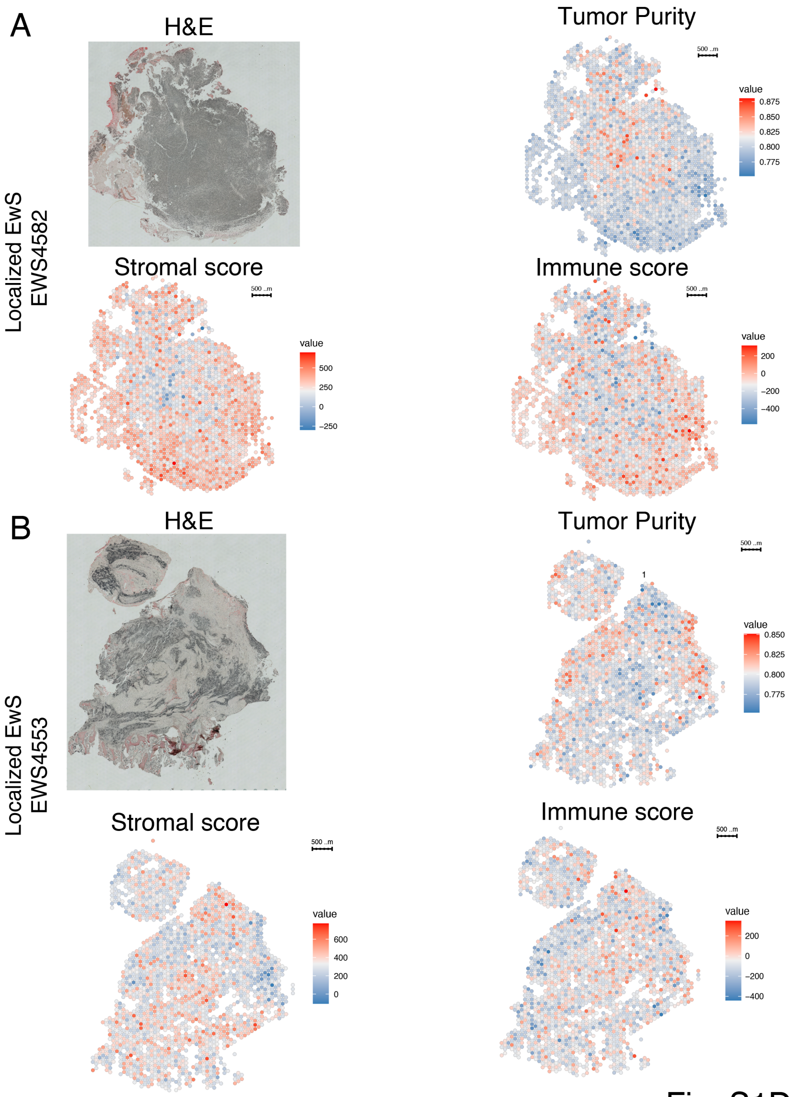

A
H&E
Tumor Purity
Localized EwS
EWS4582
Stromal score
Immune score
value
0.875
0.850
0.825
0.800
0.775
500
250
0
−250
200
0
−200
−400
B
H&E
Tumor Purity
Localized EwS
EWS4553
Stromal score
Immune score
0.850
0.825
0.800
0.775
600
400
200
0
200
0
−200
−400


Fig. S1D

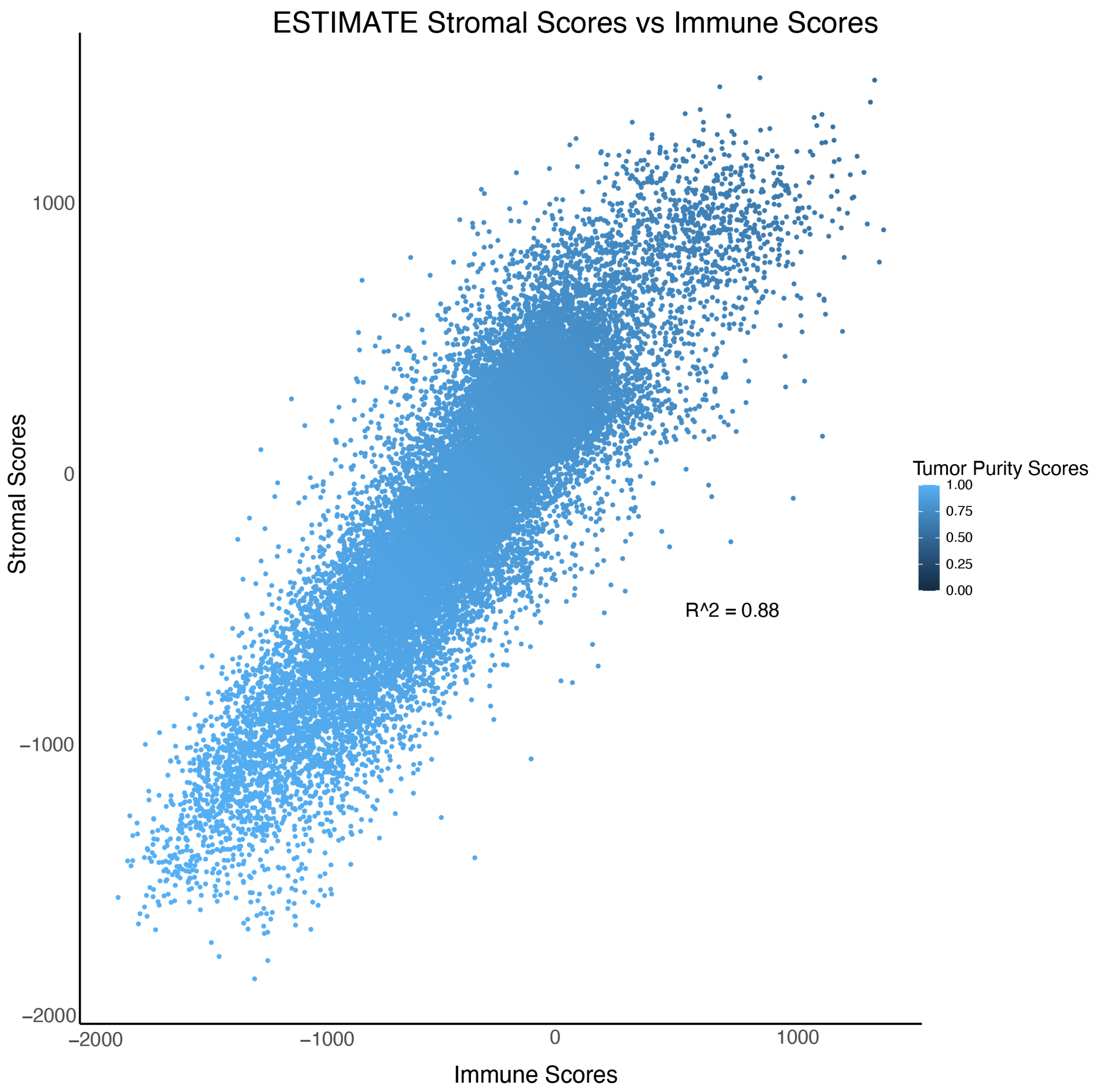

ESTIMATE Stromal Scores vs Immune Scores
Stromal Scores
1000
0
−1000
−2000
−2000
−1000
0
1000
Immune Scores
R^2 = 0.88
Tumor Purity Scores
1.00
0.75
0.50
0.25
0.00


Figure S1E

**Figure S1A-D:** ESTIMATE transcriptional analysis and representative H&E stained images of metastatic and localized EwS tumors.

**Figure S1E:** Scattered plot of immune and stromal scores per spot in all samples, with a correlation coefficient of $R^2$ = 0.88. Data points are colored by tumor purity score, with an inverse relationship to immune and stromal scores.

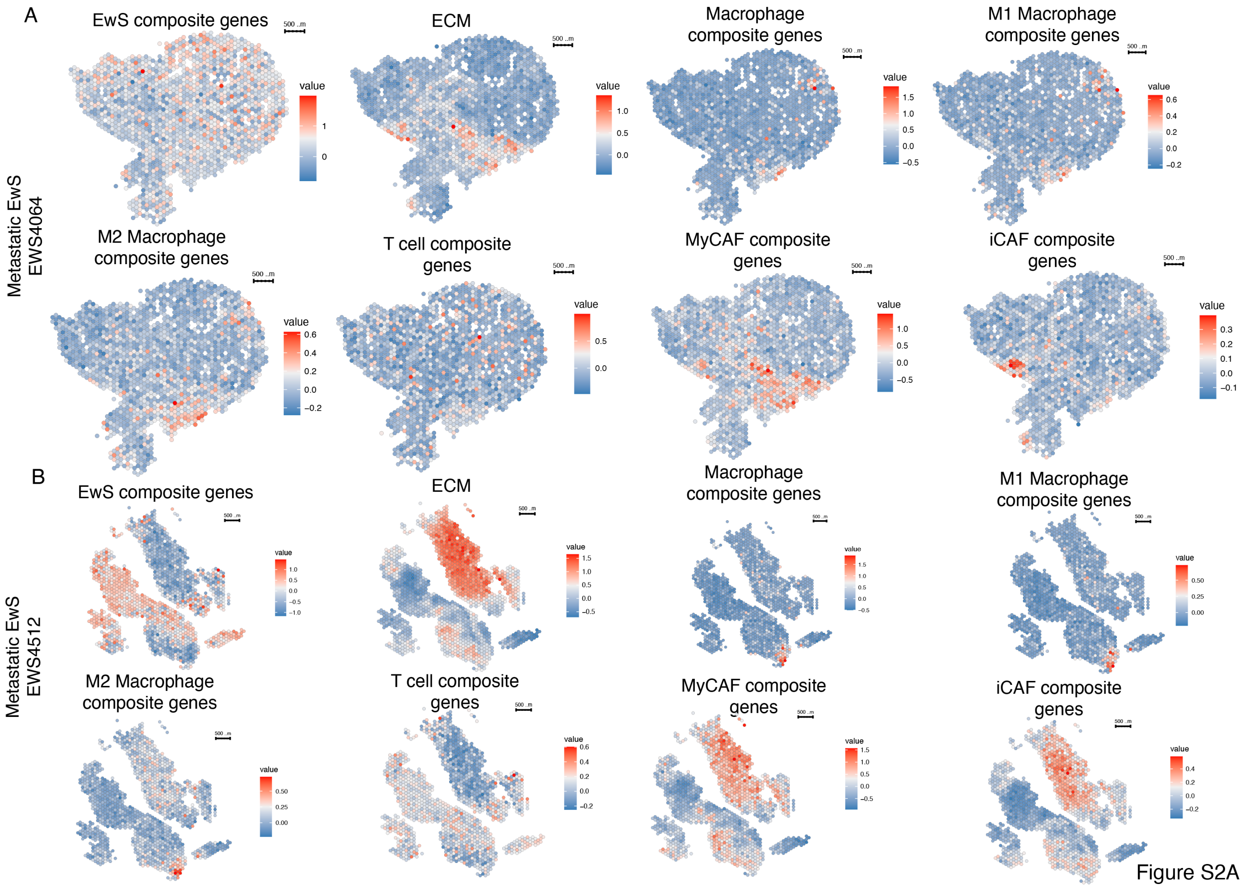
A
Metastatic EwS
EWS4064
EwS composite genes
ECM
Macrophage composite genes
M1 Macrophage composite genes
M2 Macrophage composite genes
T cell composite genes
MyCAF composite genes
iCAF composite genes
value
B
Metastatic EwS
EWS4512
EwS composite genes
ECM
Macrophage composite genes
M1 Macrophage composite genes
M2 Macrophage composite genes
T cell composite genes
MyCAF composite genes
iCAF composite genes
value

Figure S2A

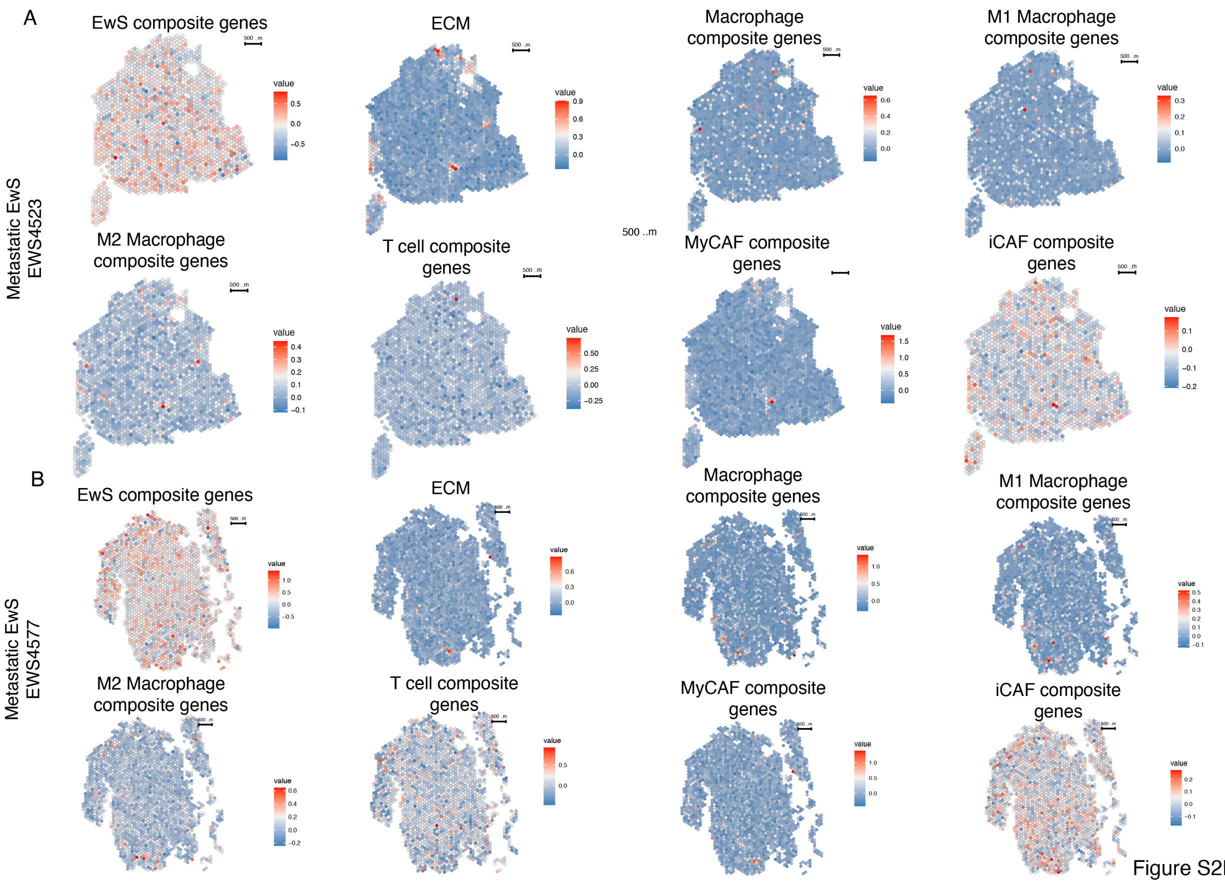
A
Metastatic EwS
EWS4523
EwS composite genes
ECM
Macrophage composite genes
M1 Macrophage composite genes
M2 Macrophage composite genes
T cell composite genes
MyCAF composite genes
iCAF composite genes
500 ..m
B
Metastatic EwS
EWS4577
EwS composite genes
ECM
Macrophage composite genes
M1 Macrophage composite genes
M2 Macrophage composite genes
T cell composite genes
MyCAF composite genes
iCAF composite genes
value

Figure S2B

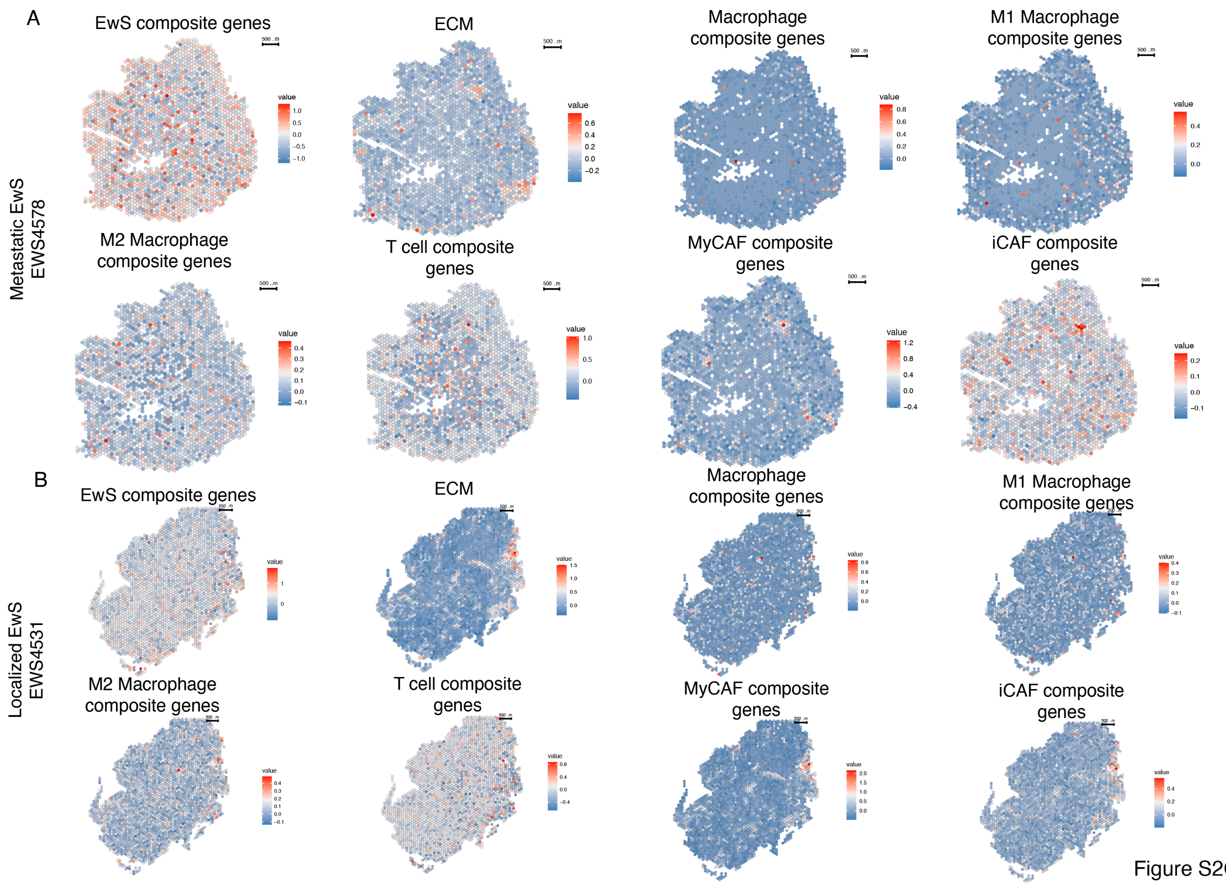
A
EwS composite genes
ECM
Macrophage composite genes
M1 Macrophage composite genes
Metastatic EwS EWS4578
M2 Macrophage composite genes
T cell composite genes
MyCAF composite genes
iCAF composite genes
B
EwS composite genes
ECM
Macrophage composite genes
M1 Macrophage composite genes
Localized EwS EWS4531
M2 Macrophage composite genes
T cell composite genes
MyCAF composite genes
iCAF composite genes

Figure S2C

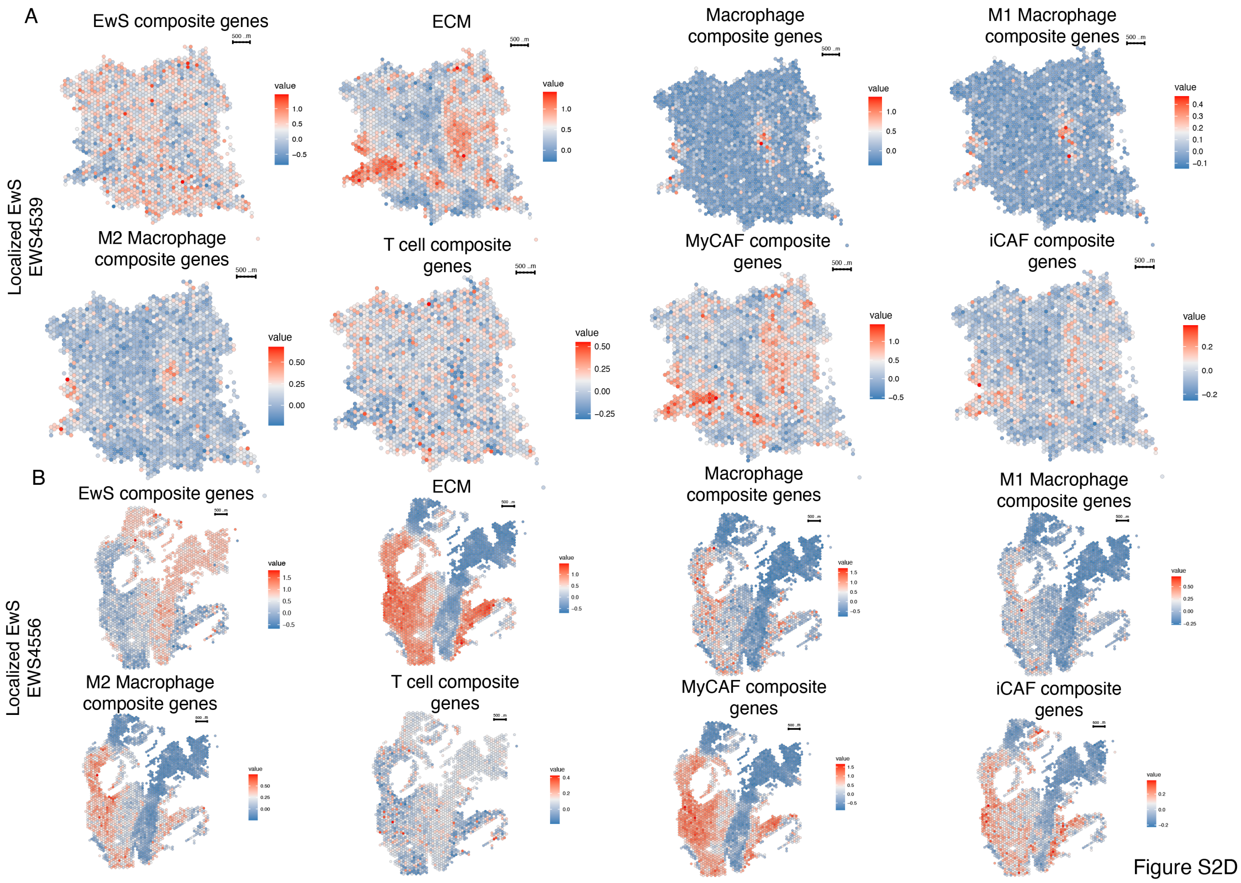
A
EwS composite genes
ECM
Macrophage composite genes
M1 Macrophage composite genes
Localized EwS EWS4539
M2 Macrophage composite genes
T cell composite genes
MyCAF composite genes
iCAF composite genes
B
Localized EwS EWS4556


Figure S2D

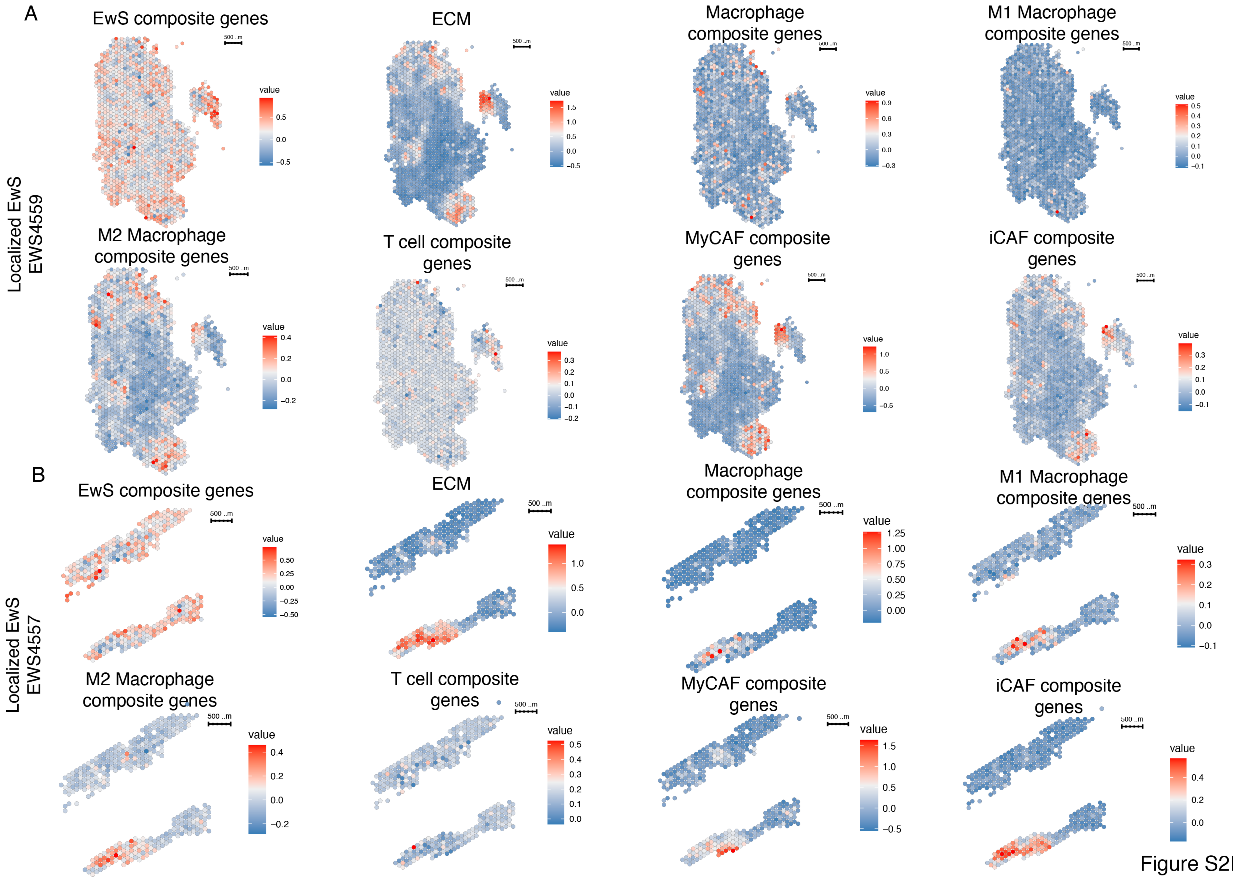
A
Localized EwS
EWS4559
EwS composite genes
ECM
Macrophage composite genes
M1 Macrophage composite genes
M2 Macrophage composite genes
T cell composite genes
MyCAF composite genes
iCAF composite genes
B
Localized EwS
EWS4557
EwS composite genes
ECM
Macrophage composite genes
M1 Macrophage composite genes
M2 Macrophage composite genes
T cell composite genes
MyCAF composite genes
iCAF composite genes
value

Figure S2E

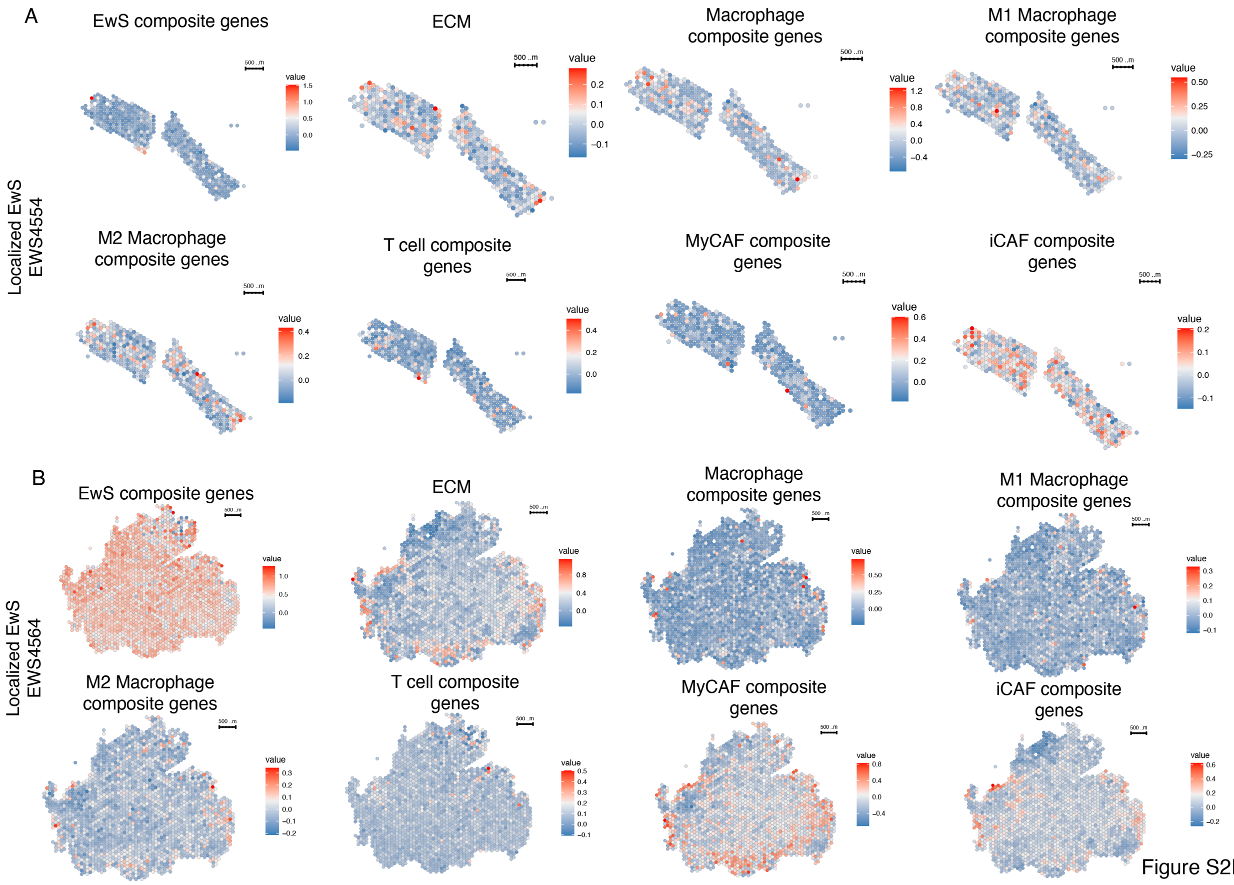
A
EwS composite genes
ECM
Macrophage composite genes
M1 Macrophage composite genes
Localized EwS EWS4554
M2 Macrophage composite genes
T cell composite genes
MyCAF composite genes
iCAF composite genes
B
EwS composite genes
ECM
Macrophage composite genes
M1 Macrophage composite genes
Localized EwS EWS4564
M2 Macrophage composite genes
T cell composite genes
MyCAF composite genes
iCAF composite genes
value

Figure S2F

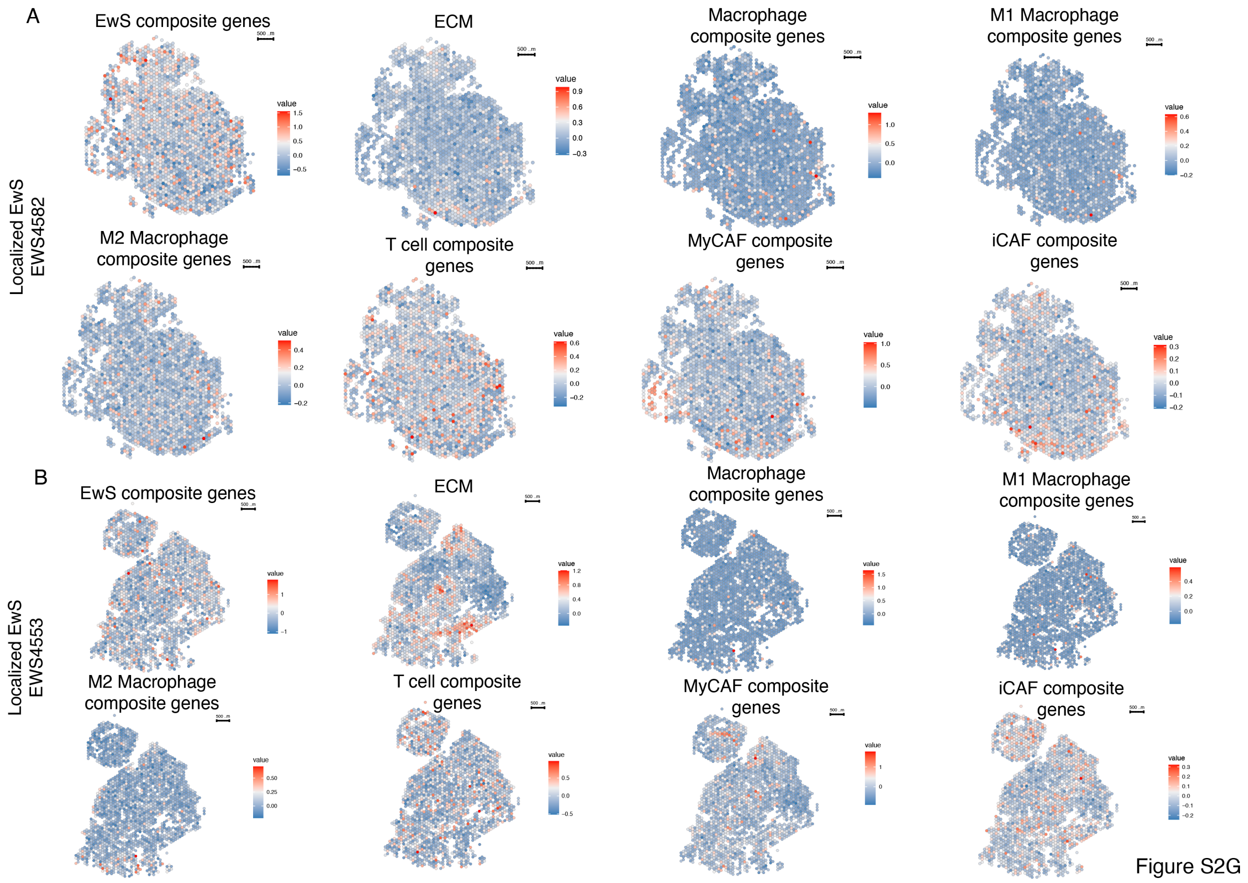


Figure S2A-G: Spatial gene enrichment mapping of composite genes annotating EwS, ECM, and immune cells in both localized and metastatic EwS tumors.

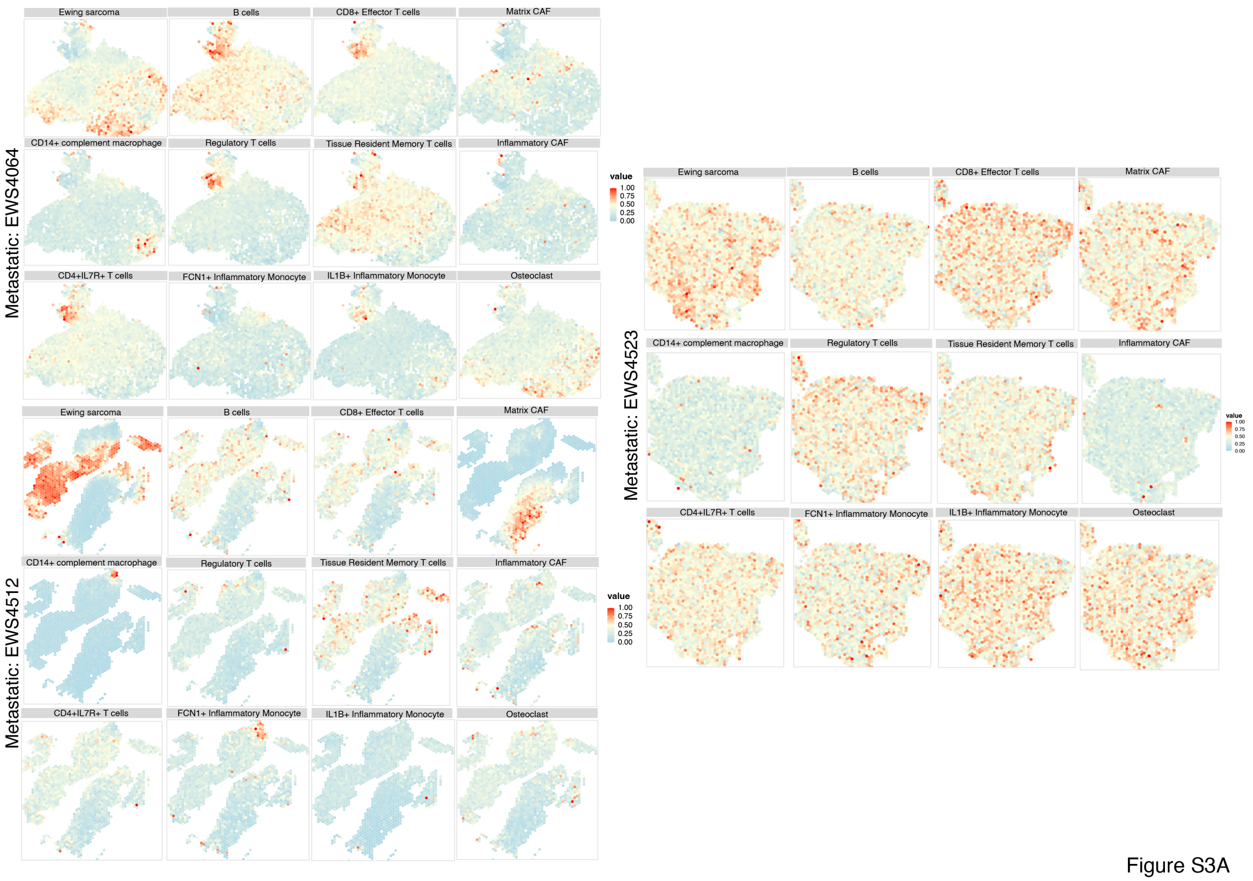
Metastatic: EWS4064
Ewing sarcoma
B cells
CD8+ Effector T cells
Matrix CAF
CD14+ complement macrophage
Regulatory T cells
Tissue Resident Memory T cells
Inflammatory CAF
CD4+IL7R+ T cells
FCN1+ Inflammatory Monocyte
IL1B+ Inflammatory Monocyte
Osteoclast
value
1.00
0.75
0.50
0.25
0.00
Metastatic: EWS4512
Ewing sarcoma
B cells
CD8+ Effector T cells
Matrix CAF
CD14+ complement macrophage
Regulatory T cells
Tissue Resident Memory T cells
Inflammatory CAF
CD4+IL7R+ T cells
FCN1+ Inflammatory Monocyte
IL1B+ Inflammatory Monocyte
Osteoclast
value
1.00
0.75
0.50
0.25
0.00
Metastatic: EWS4523
Ewing sarcoma
B cells
CD8+ Effector T cells
Matrix CAF
CD14+ complement macrophage
Regulatory T cells
Tissue Resident Memory T cells
Inflammatory CAF
CD4+IL7R+ T cells
FCN1+ Inflammatory Monocyte
IL1B+ Inflammatory Monocyte
Osteoclast

Figure S3A

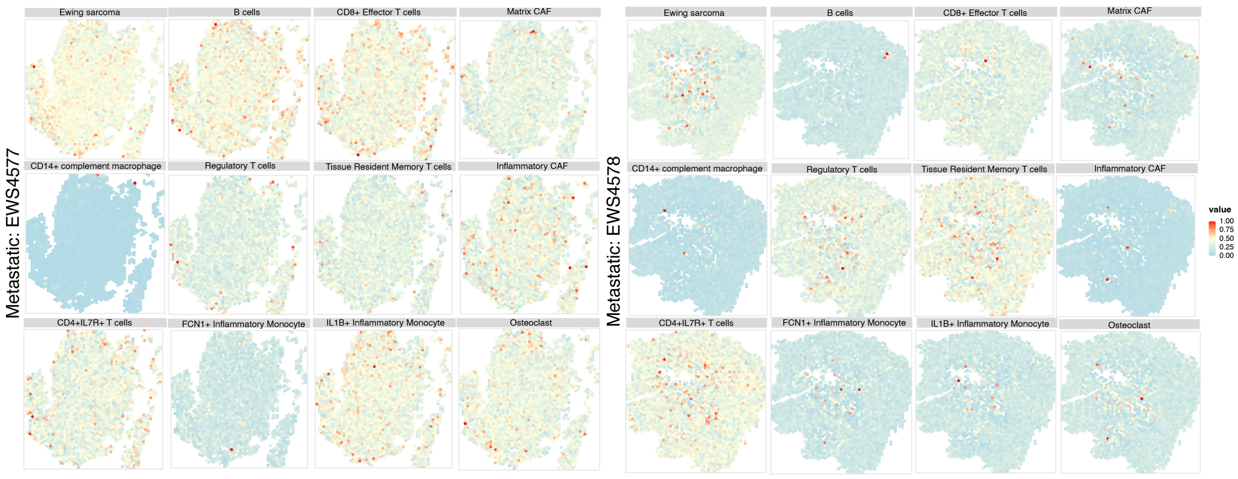
Metastatic: EWS4577
Ewing sarcoma
B cells
CD8+ Effector T cells
Matrix CAF
CD14+ complement macrophage
Regulatory T cells
Tissue Resident Memory T cells
Inflammatory CAF
CD4+IL7R+ T cells
FCN1+ Inflammatory Monocyte
IL1B+ Inflammatory Monocyte
Osteoclast
Metastatic: EWS4578
Ewing sarcoma
B cells
CD8+ Effector T cells
Matrix CAF
CD14+ complement macrophage
Regulatory T cells
Tissue Resident Memory T cells
Inflammatory CAF
CD4+IL7R+ T cells
FCN1+ Inflammatory Monocyte
IL1B+ Inflammatory Monocyte
Osteoclast
value
1.00
0.75
0.50
0.25
0.00

Figure S3B

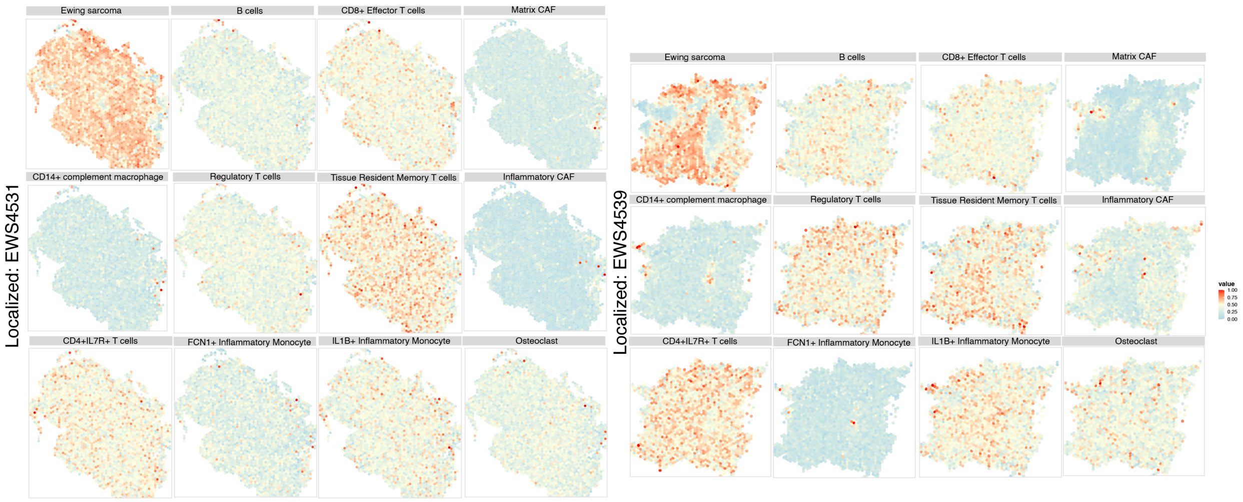
Localized: EWS4531
Ewing sarcoma
B cells
CD8+ Effector T cells
Matrix CAF
CD14+ complement macrophage
Regulatory T cells
Tissue Resident Memory T cells
Inflammatory CAF
CD4+IL7R+ T cells
FCN1+ Inflammatory Monocyte
IL1B+ Inflammatory Monocyte
Osteoclast
Localized: EWS4539
Ewing sarcoma
B cells
CD8+ Effector T cells
Matrix CAF
CD14+ complement macrophage
Regulatory T cells
Tissue Resident Memory T cells
Inflammatory CAF
CD4+IL7R+ T cells
FCN1+ Inflammatory Monocyte
IL1B+ Inflammatory Monocyte
Osteoclast
value
1.00
0.75
0.50
0.25
0.00

Figure S3C

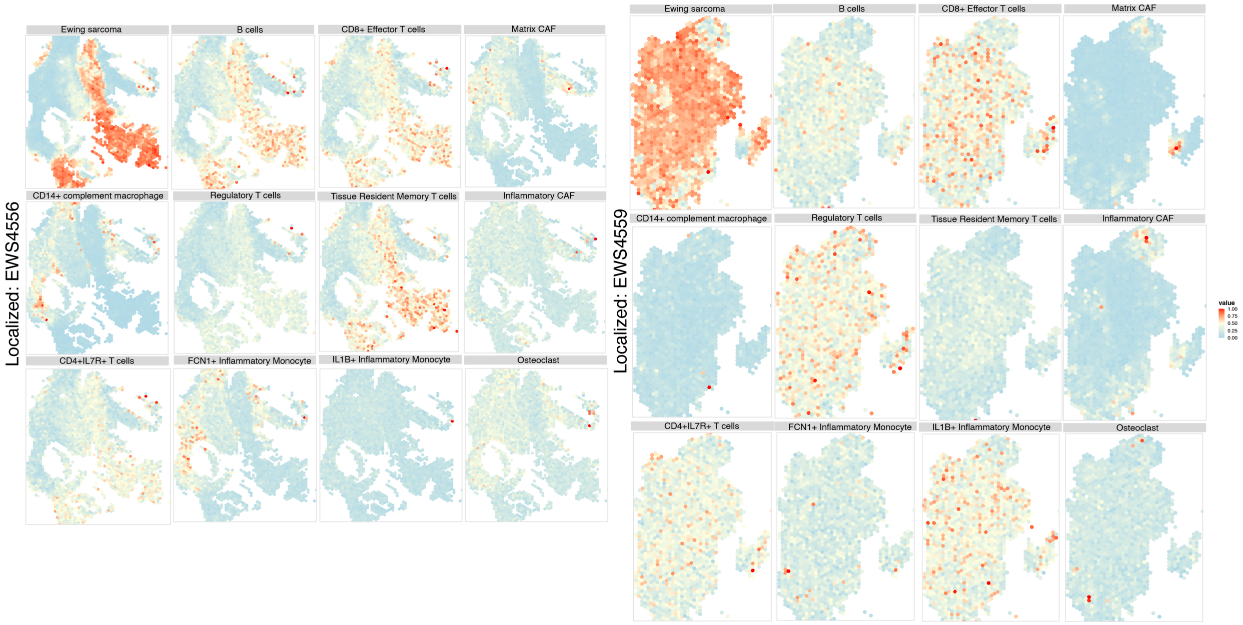
Localized: EWS4556
Ewing sarcoma
B cells
CD8+ Effector T cells
Matrix CAF
CD14+ complement macrophage
Regulatory T cells
Tissue Resident Memory T cells
Inflammatory CAF
CD4+IL7R+ T cells
FCN1+ Inflammatory Monocyte
IL1B+ Inflammatory Monocyte
Osteoclast
Localized: EWS4559
Ewing sarcoma
B cells
CD8+ Effector T cells
Matrix CAF
CD14+ complement macrophage
Regulatory T cells
Tissue Resident Memory T cells
Inflammatory CAF
CD4+IL7R+ T cells
FCN1+ Inflammatory Monocyte
IL1B+ Inflammatory Monocyte
Osteoclast
value

Figure S3D

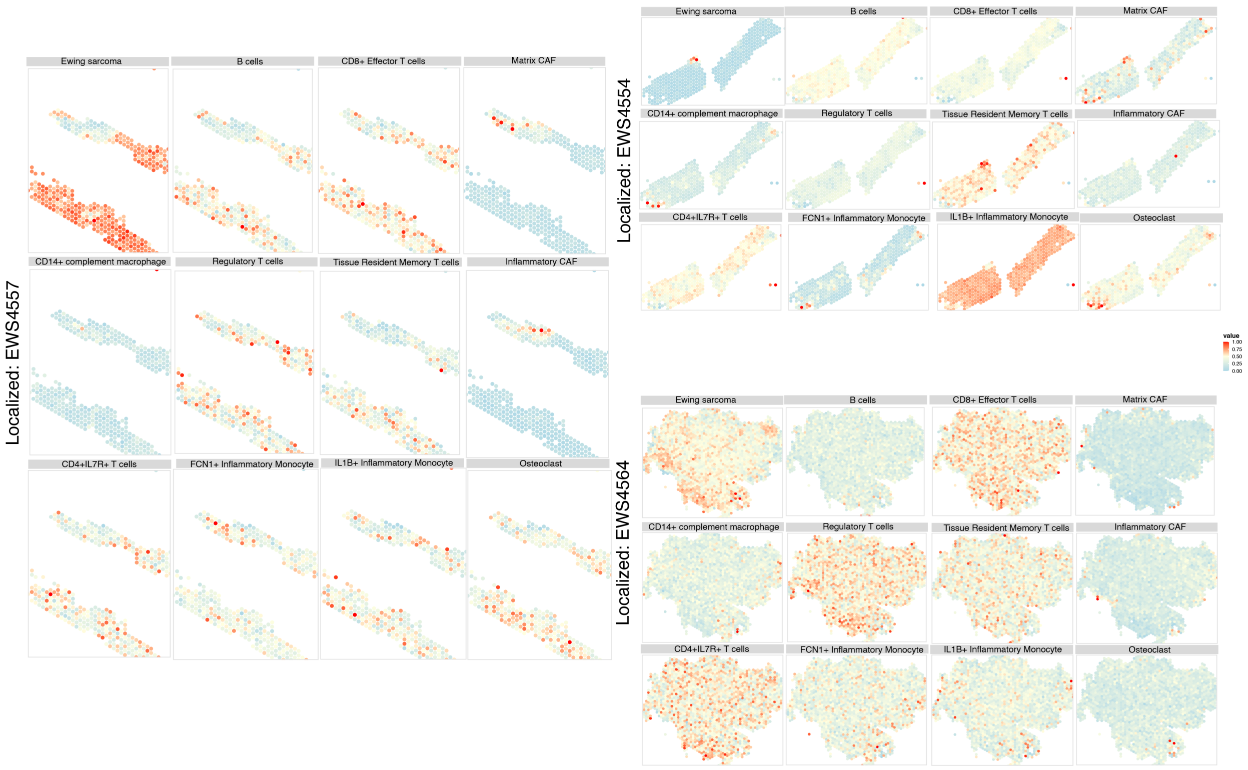
Localized: EWS4557
Ewing sarcoma
B cells
CD8+ Effector T cells
Matrix CAF
CD14+ complement macrophage
Regulatory T cells
Tissue Resident Memory T cells
Inflammatory CAF
CD4+IL7R+ T cells
FCN1+ Inflammatory Monocyte
IL1B+ Inflammatory Monocyte
Osteoclast
Localized: EWS4554
Localized: EWS4564
value
1.00
0.75
0.50
0.25
0.00

Figure S3E

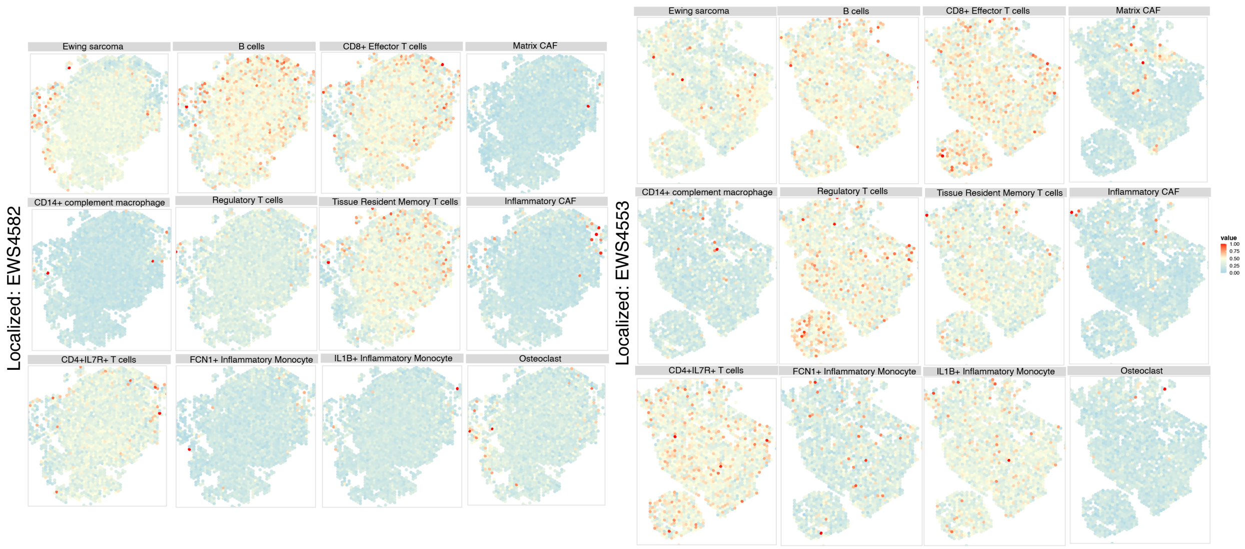


Figure S3F

**Figure S3A-F:** Spatial visualization of deconvolution analysis of localized and metastatic EwS tumors.

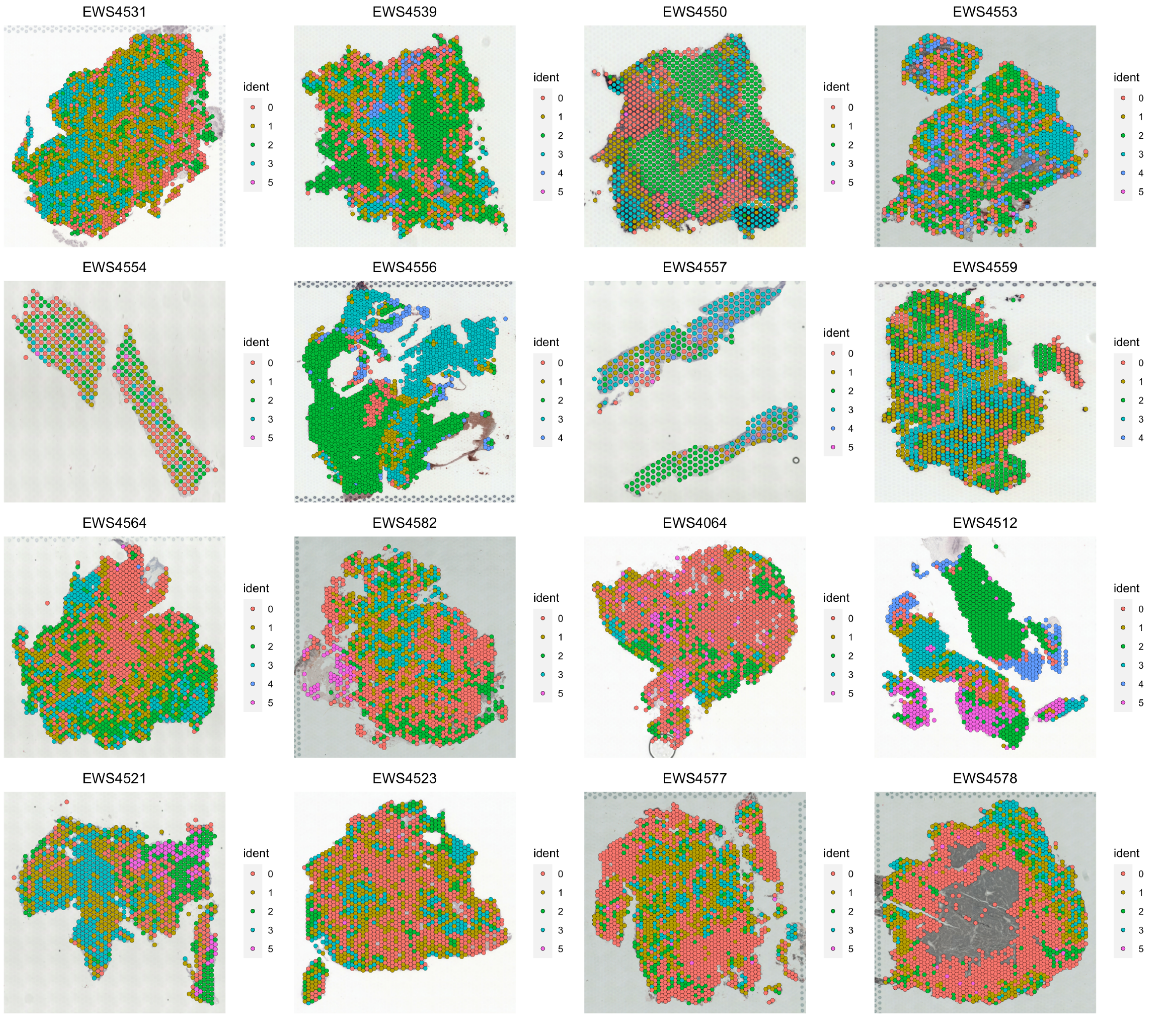


**Figure S4:** Spatial plotting of integrated transcriptomic phenotypic clustering of all 16 tumors prior to NICHES analysis.

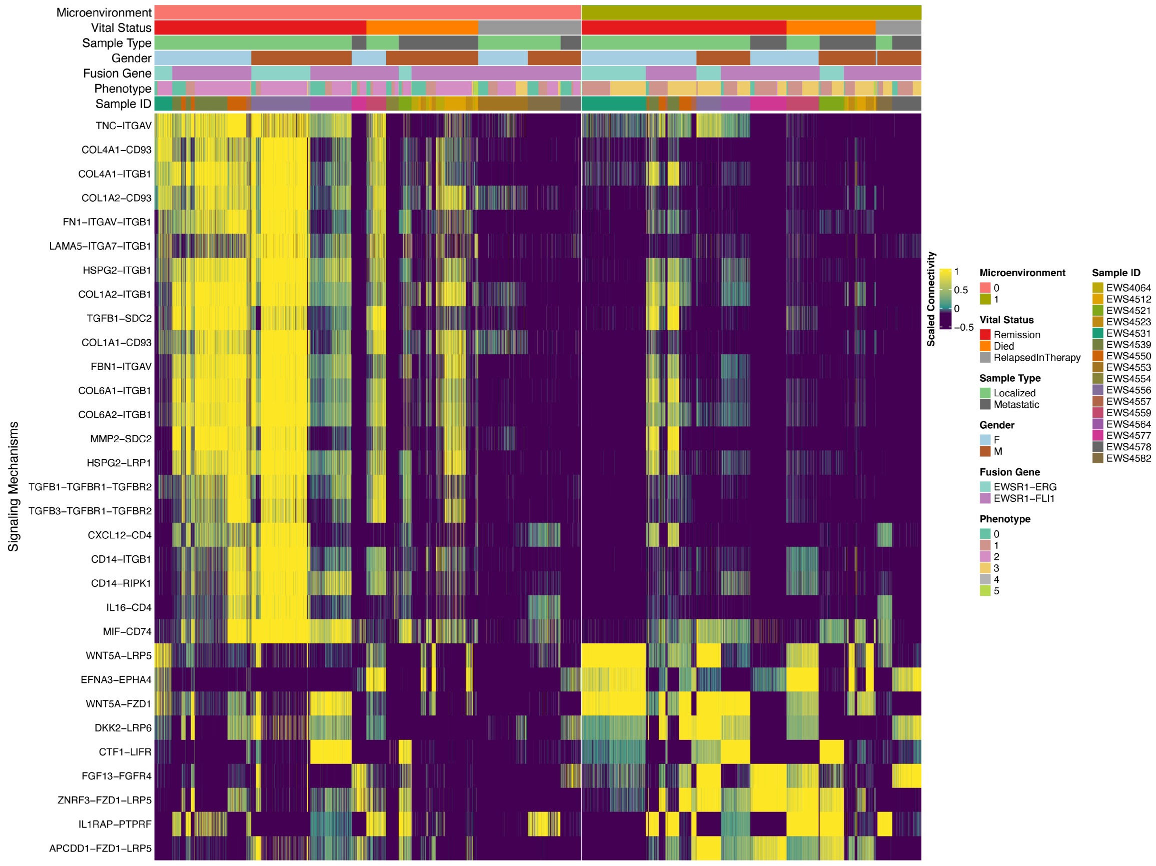


Figure S5

**Figure S5:** Complex heatmap of NICHES analysis with microenvironmental connectivity clusters, vital status, sample type, gender, fusion gene, phenotype and sample ID.

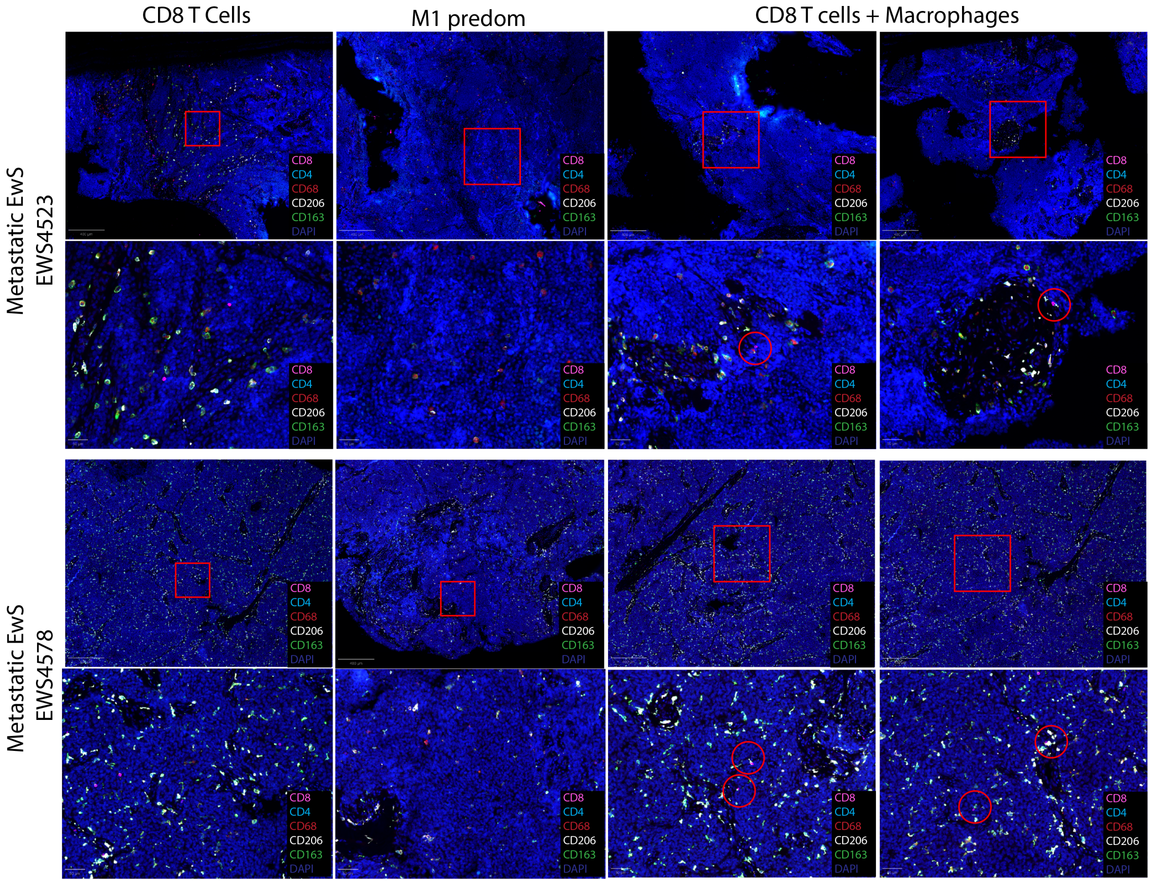
CD8 T Cells
M1 predom
CD8 T cells + Macrophages
Metastatic EwS
EWS4523
Metastatic EwS
EWS4578
CD8
CD4
CD68
CD206
CD163
DAPI

Figure S6A

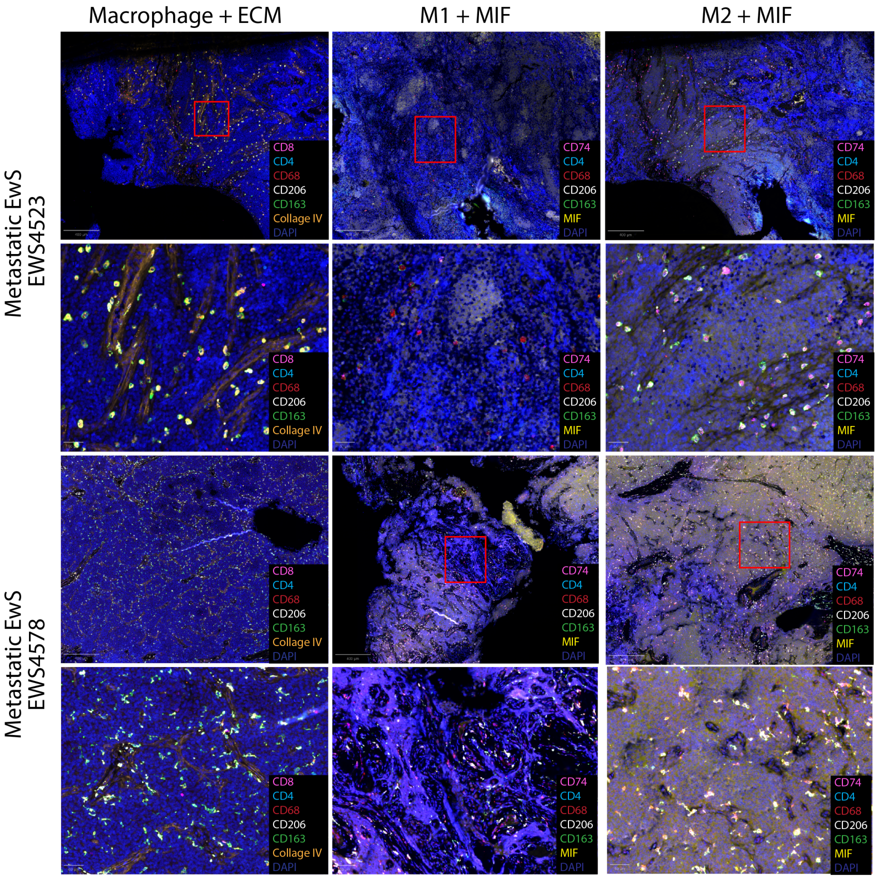

Macrophage + ECM
M1 + MIF
M2 + MIF
Metastatic EwS EWS4523
Metastatic EwS EWS4578
CD8
CD4
CD68
CD206
CD163
Collage IV
DAPI
CD74
MIF


Figure S6B

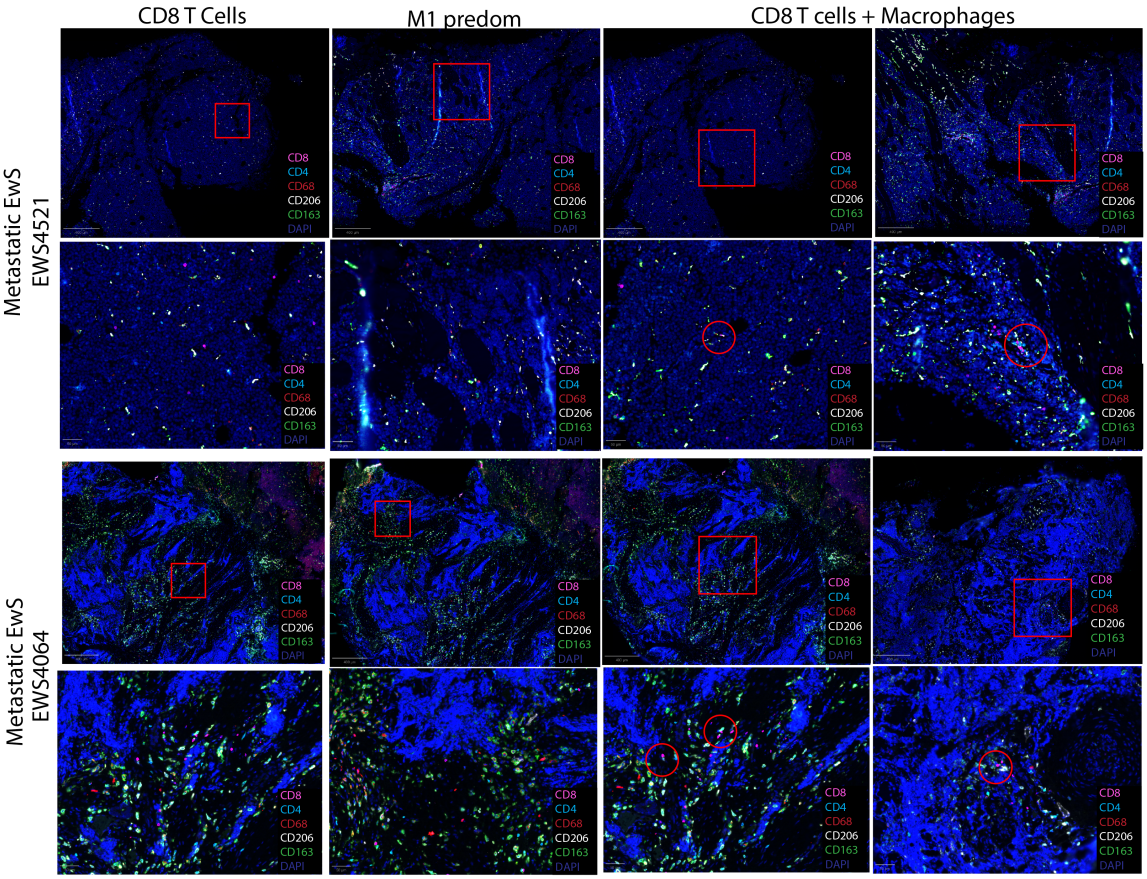
CD8 T Cells
M1 predom
CD8 T cells + Macrophages
Metastatic EwS EWS4521
Metastatic EwS EWS4064
CD8
CD4
CD68
CD206
CD163
DAPI

Figure S6C

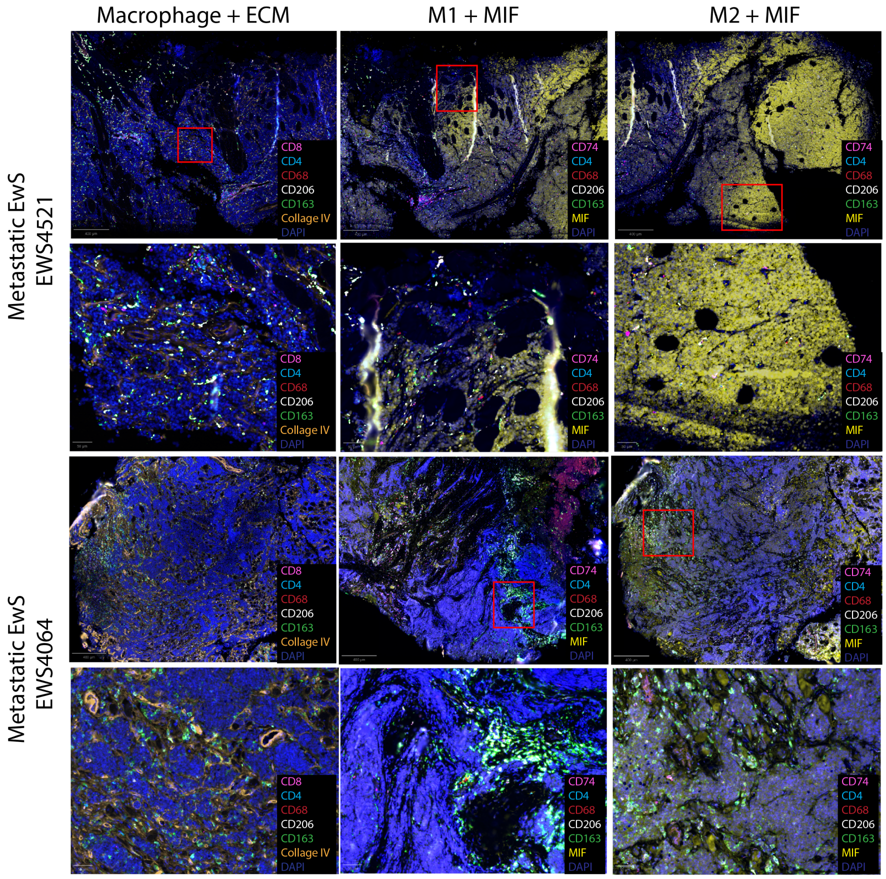
Macrophage + ECM
M1 + MIF
M2 + MIF
Metastatic EwS EWS4521
Metastatic EwS EWS4064
CD8
CD4
CD68
CD206
CD163
Collage IV
DAPI
CD74
CD4
CD68
CD206
CD163
MIF
DAPI

Figure S6D

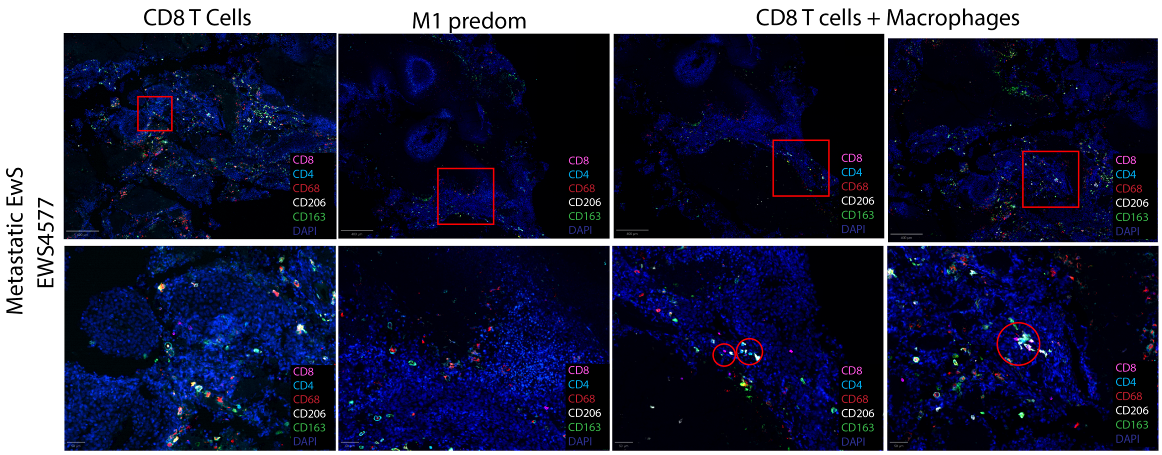
CD8 T Cells
M1 predom
CD8 T cells + Macrophages
Metastatic EwS
EWS4577
CD8
CD4
CD68
CD206
CD163
DAPI

Figure S6E

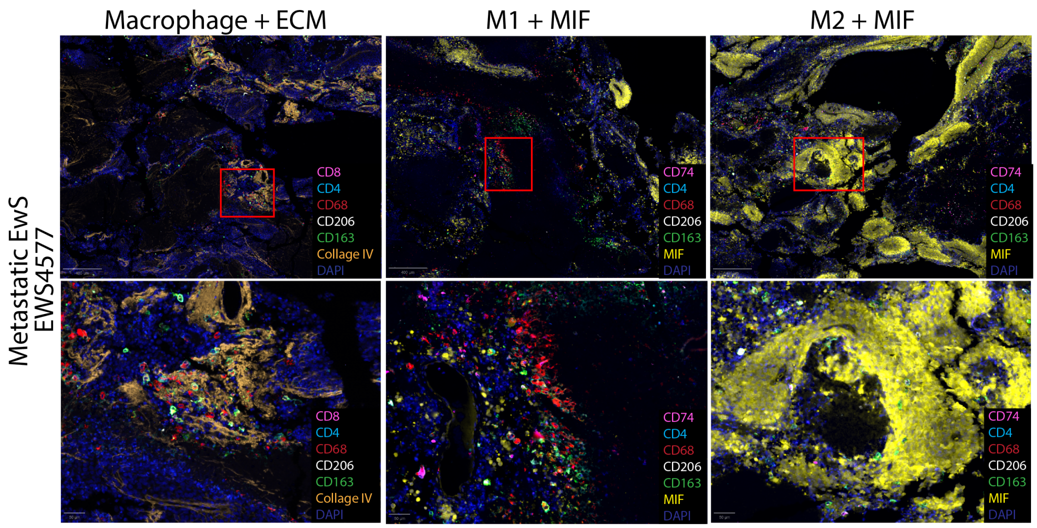

Macrophage + ECM
M1 + MIF
M2 + MIF
Metastatic EwS
EWS4577
CD8
CD4
CD68
CD206
CD163
Collage IV
DAPI
CD74
CD4
CD68
CD206
CD163
MIF
DAPI


Figure S6F

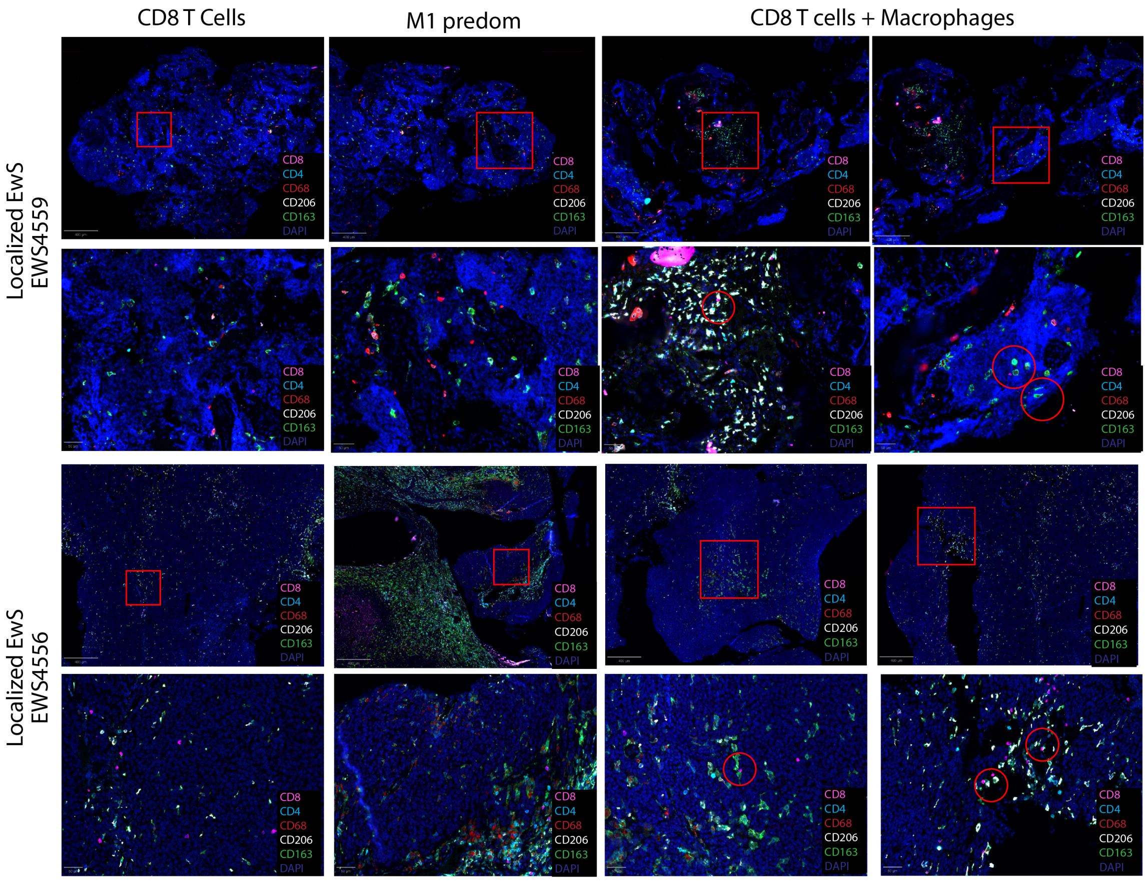
CD8 T Cells
M1 predom
CD8 T cells + Macrophages
Localized EwS EWS4559
Localized EwS EWS4556
CD8
CD4
CD68
CD206
CD163
DAPI

Figure S6G

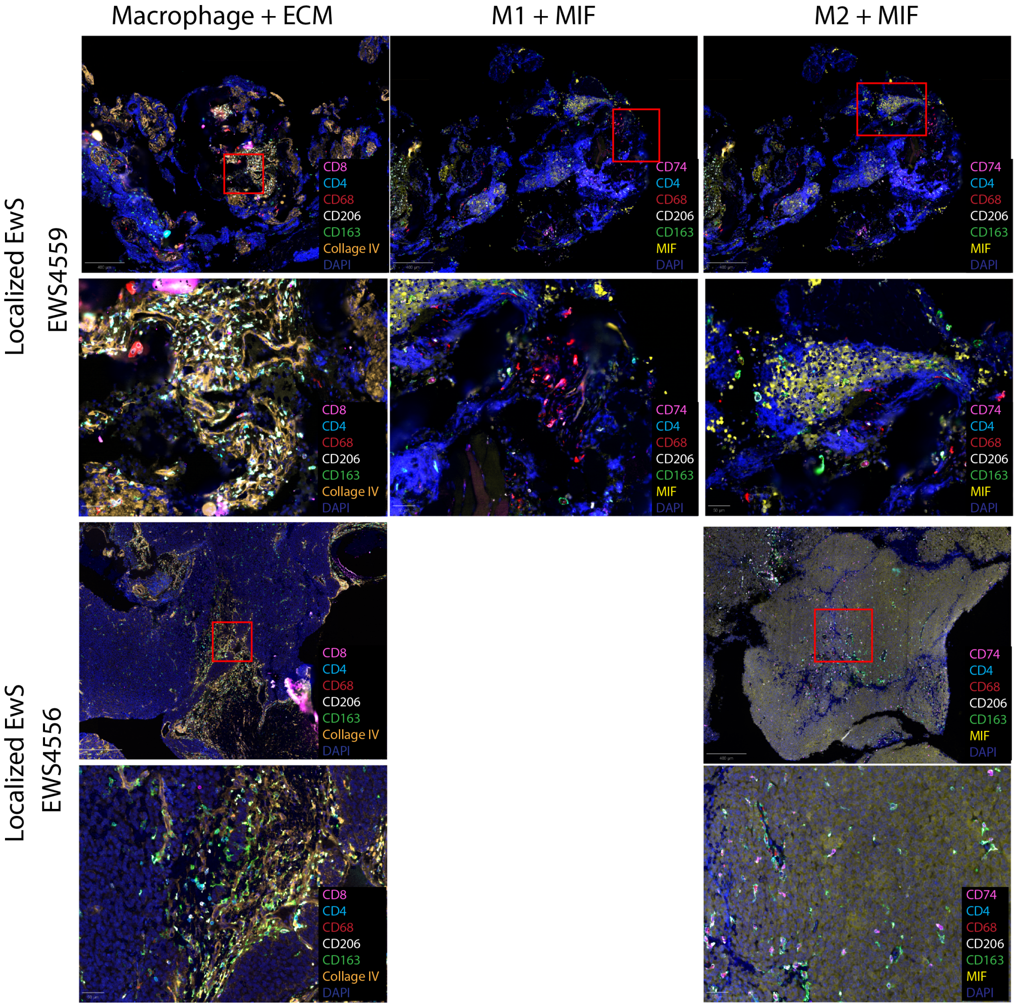
Macrophage + ECM
M1 + MIF
M2 + MIF
Localized EwS
EWS4559
Localized EwS
EWS4556
CD8
CD4
CD68
CD206
CD163
Collage IV
DAPI
CD74
CD4
CD68
CD206
CD163
MIF
DAPI

Figure S6H

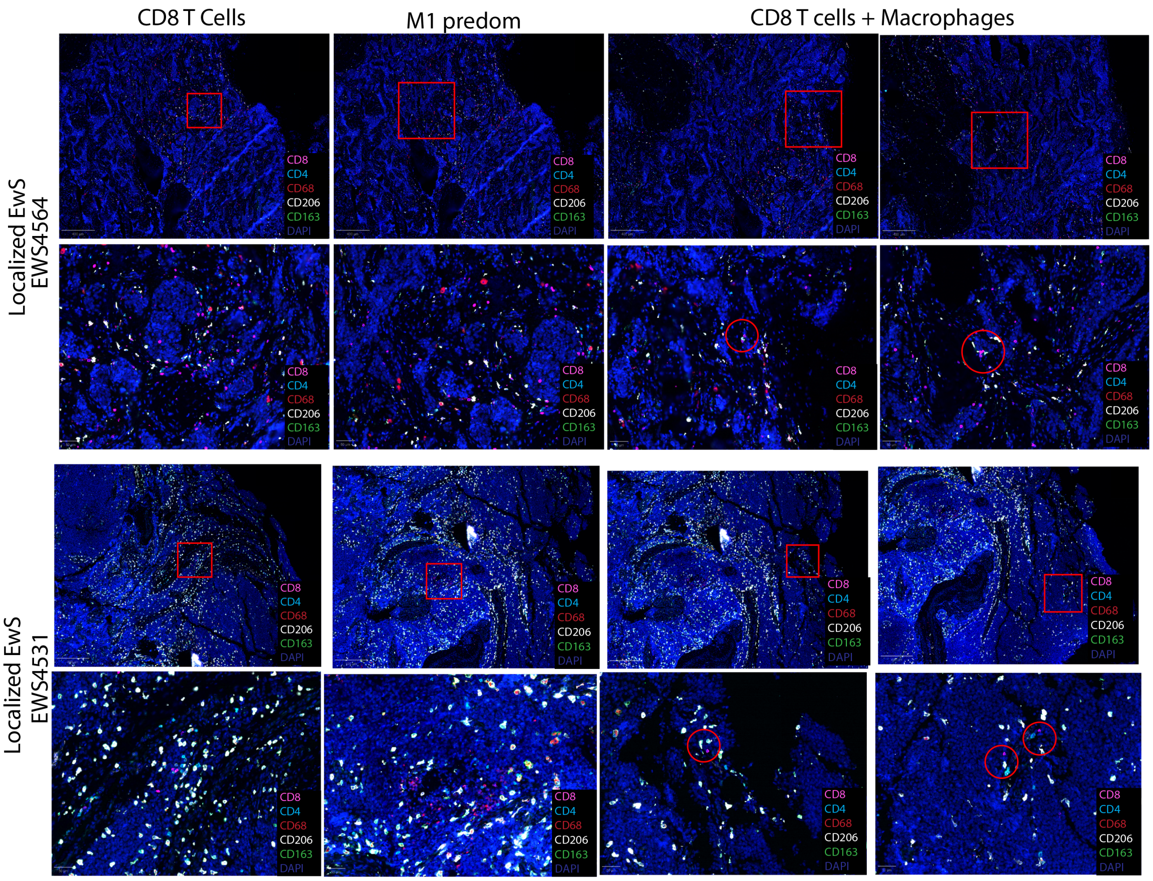
CD8 T Cells
M1 predom
CD8 T cells + Macrophages
Localized EwS
EWS4564
Localized EwS
EWS4531
CD8
CD4
CD68
CD206
CD163
DAPI

Figure S6I

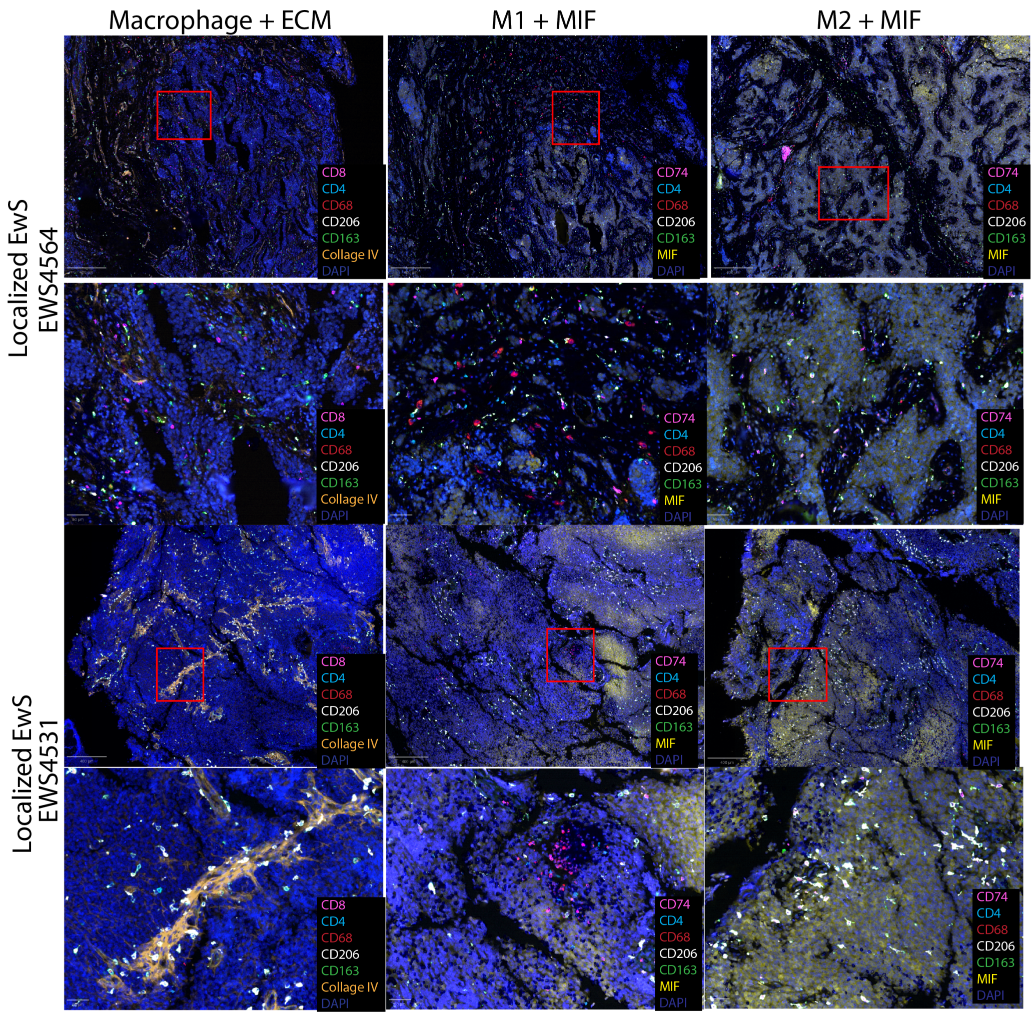

Macrophage + ECM
M1 + MIF
M2 + MIF
Localized EwS
EWS4564
Localized EwS
EWS4531
CD8
CD4
CD68
CD206
CD163
Collage IV
DAPI
CD74
MIF


Figure S6J

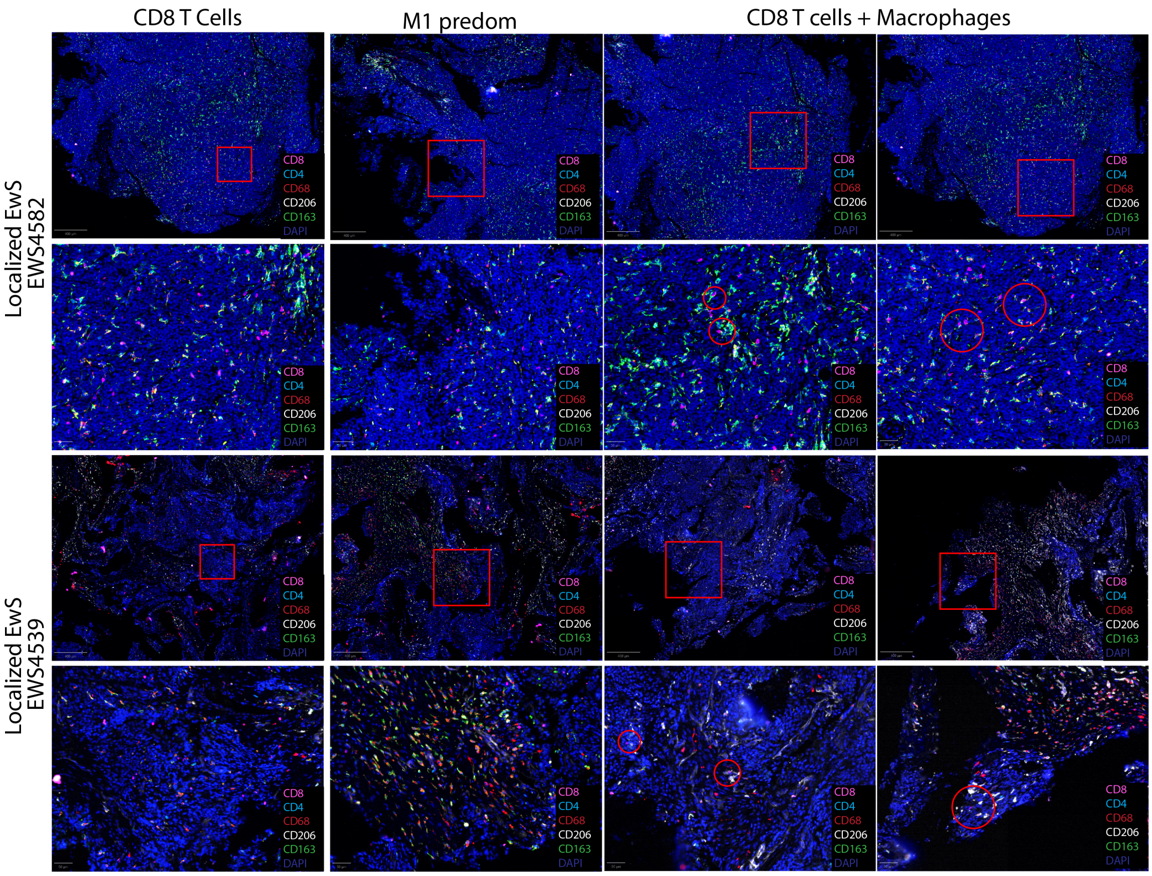

CD8 T Cells
M1 predom
CD8 T cells + Macrophages
Localized EwS
EWS4582
Localized EwS
EWS4539
CD8
CD4
CD68
CD206
CD163
DAPI


Figure S6K

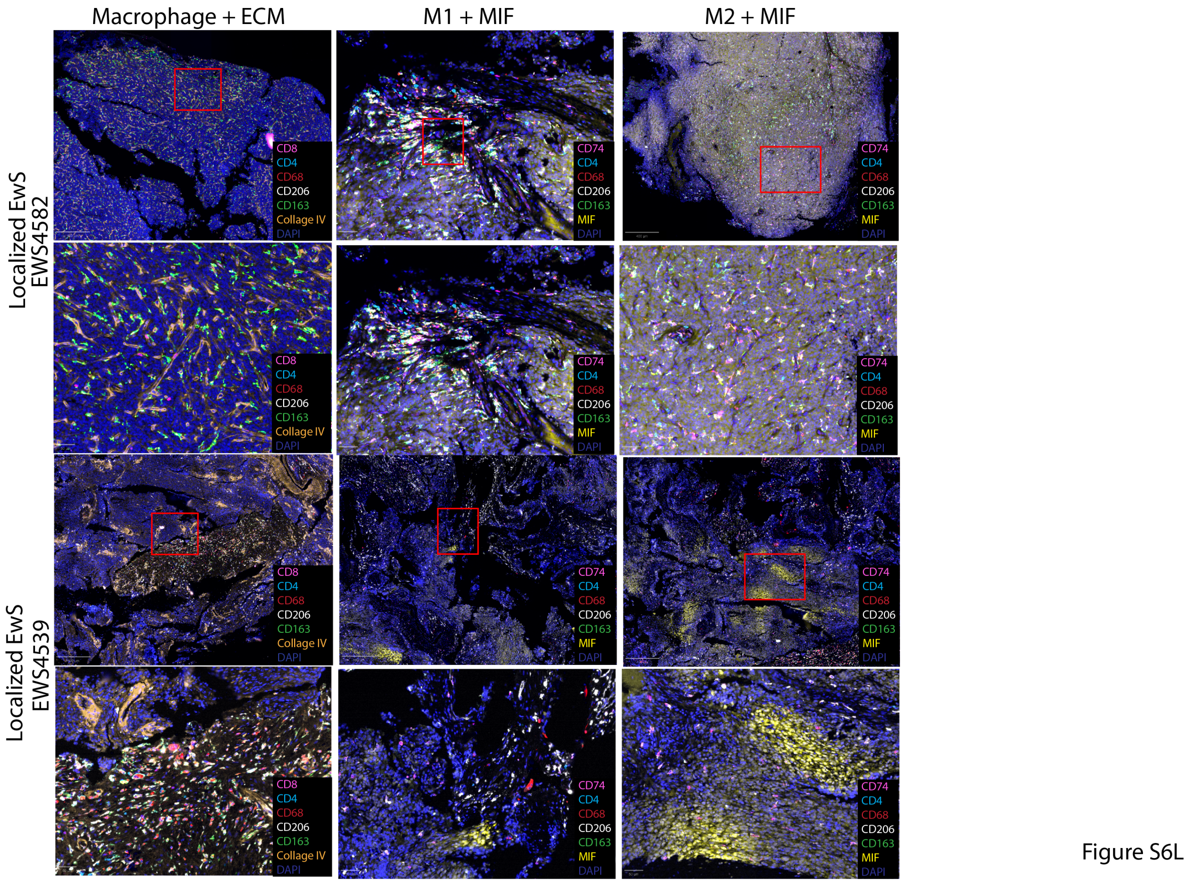


**Figure S6A-L:** Ultrahigh-plex single cell spatial proteomic analyses of localized and metastatic EwS tumors.

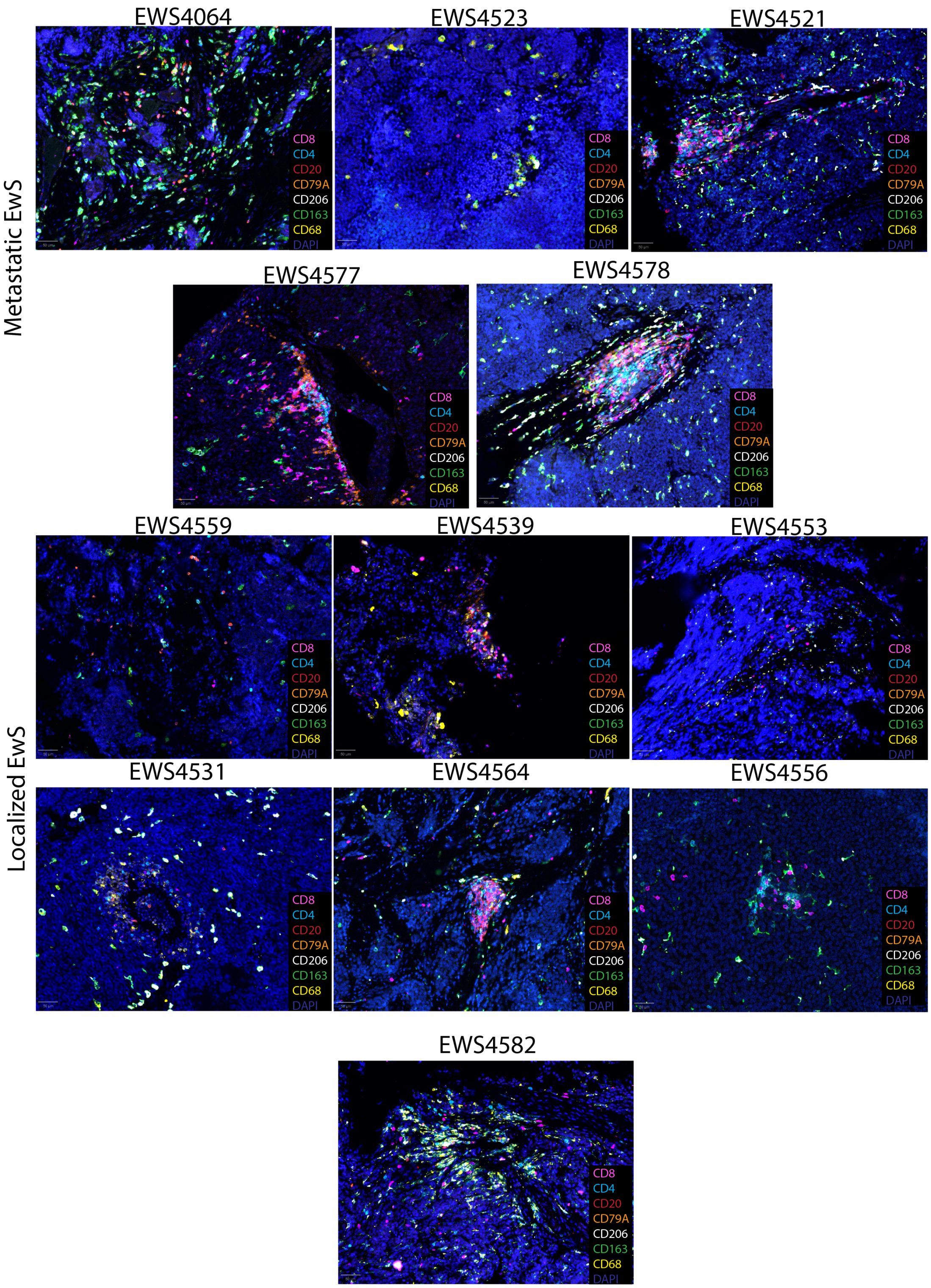


Figure S7

**Figure S7:** Ultrahigh-plex single cell spatial proteomic analyses of lymphoid aggregates discovered in localized and metastatic EwS tumors.

Met-EwS Tumors

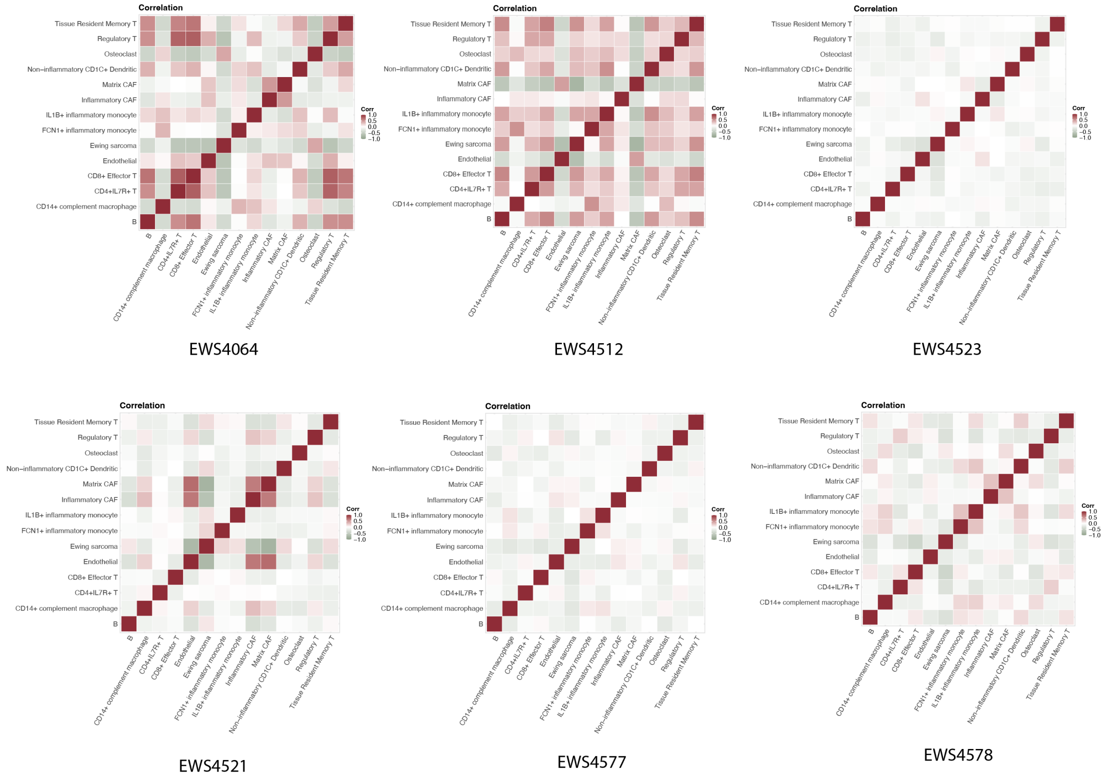


FIgure S8A

### Loc-EwS Tumors

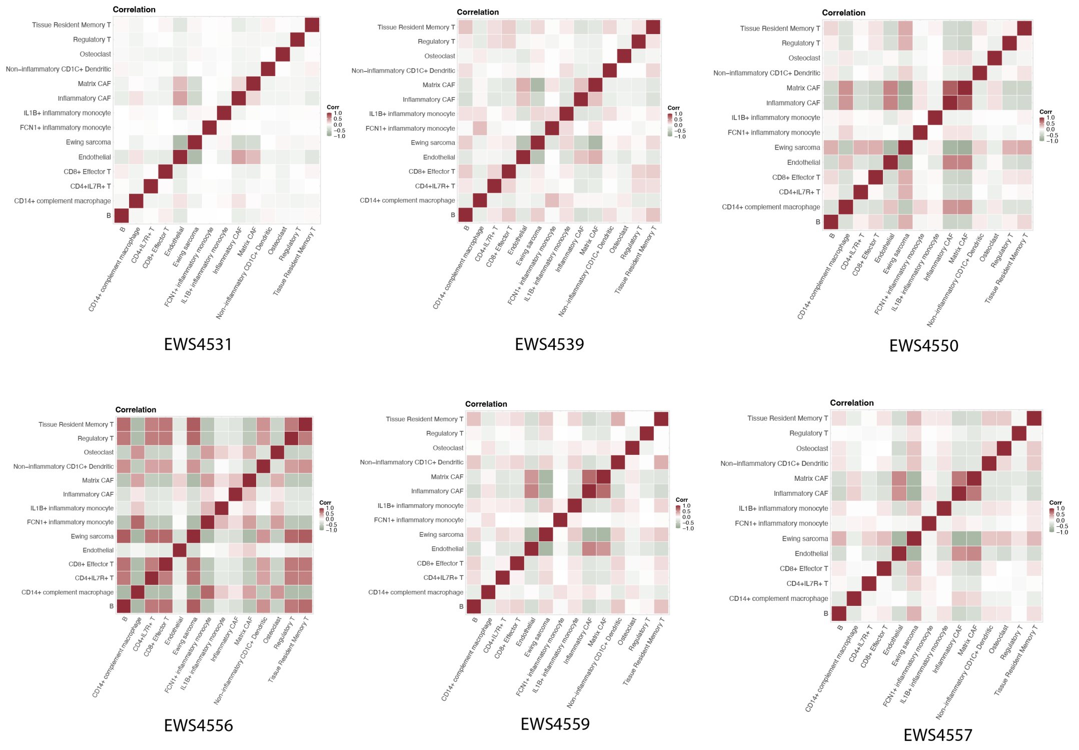


Figure S8B

Loc-EwS Tumors

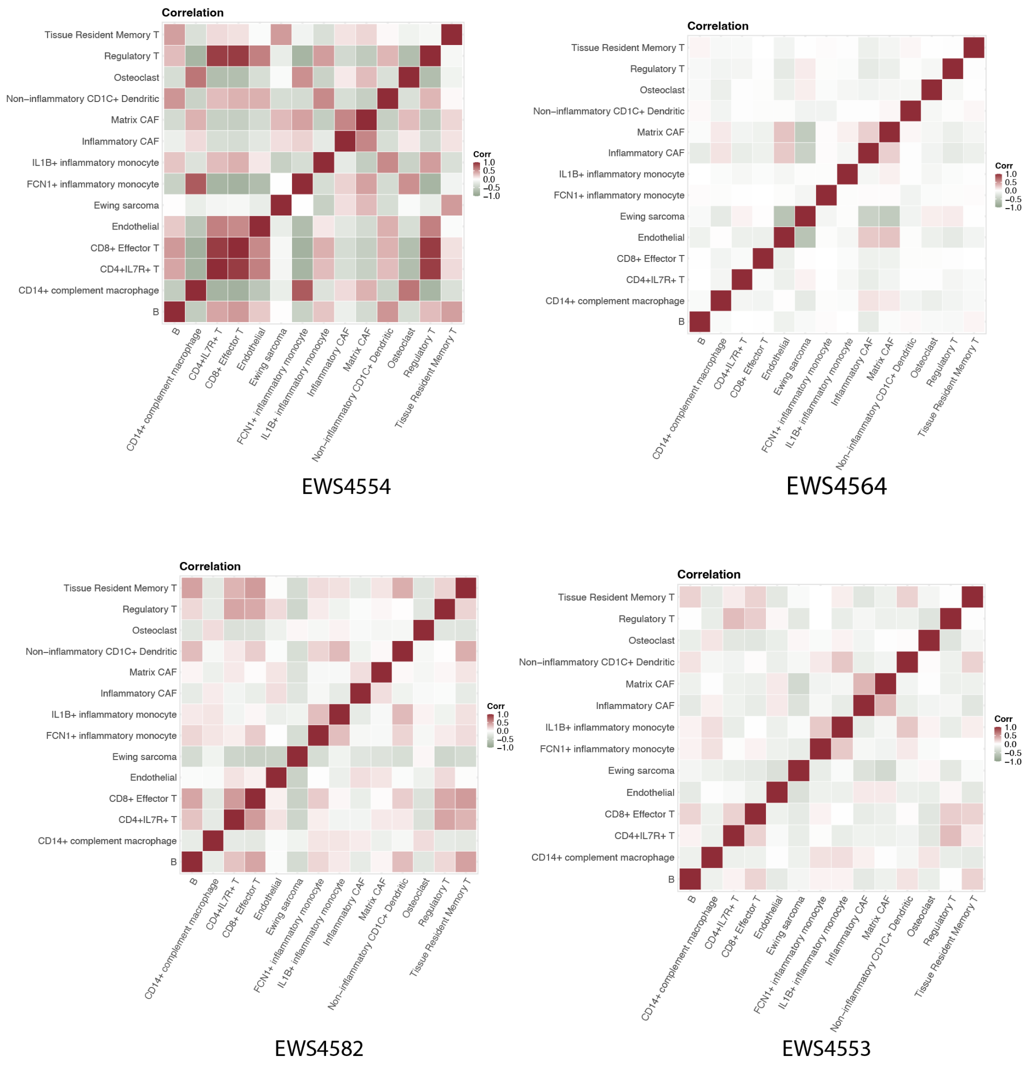


Figure S8C

**Figure S8A-C:** Correlation heatmap with deconvoluted immune cell types in both localized and metastatic EwS tumors.

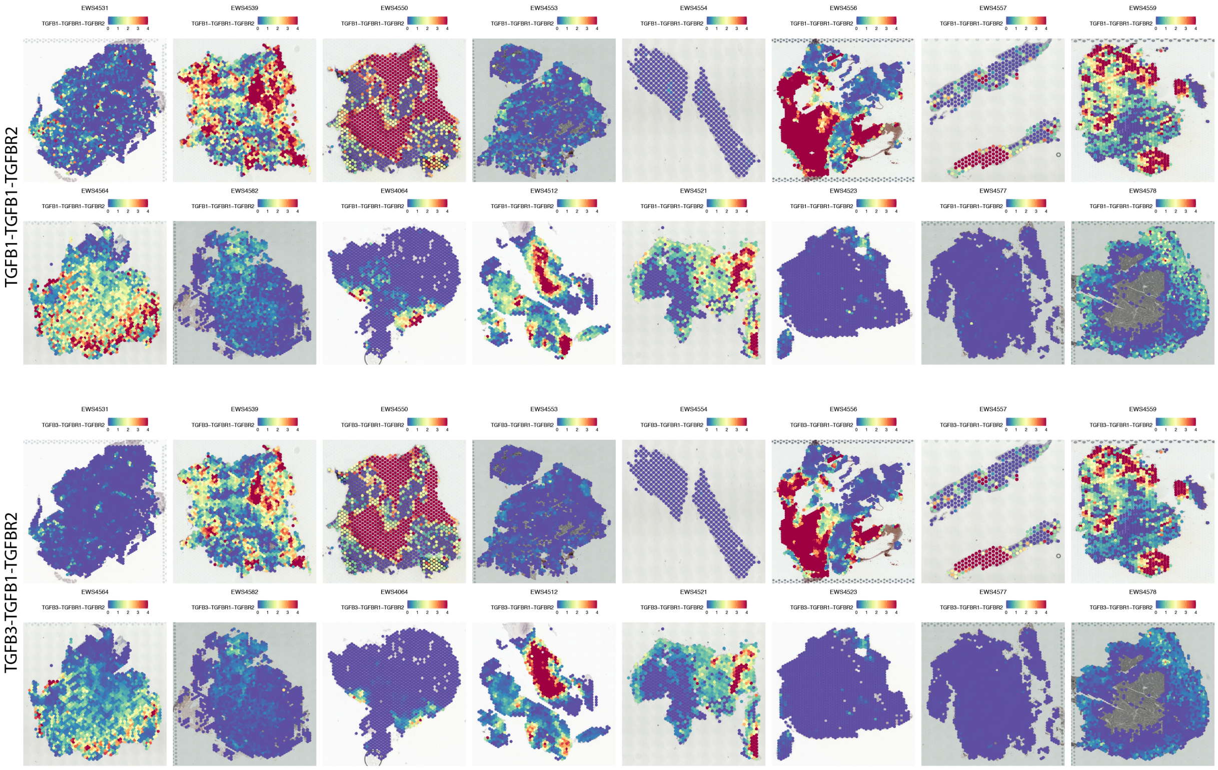


Figure S9

**Figure S9**: NICHES spatial mapping of *TGFB1-TGFBR1-TGFBR2* and *TGFB3-TGFBR1-TGFBR2* in localized and metastatic EwS tumors.
